# Multi-point convective delivery overcomes mass transport barriers for myocardial therapeutics

**DOI:** 10.64898/2026.04.29.721610

**Authors:** Jihoon Park, Adwik Rahematpura, Eva Beresin, Aaditya Majumdar, Yusra Azeem, Haruto Mizukai, Ramy Ghanim, Joy Jackson, Sean Healy, Julia Z. Ding, McKenna Clinch, Abdulraouf M. Abbas, Megan Belanger, James E. Dahlman, Joshua L. Chan, Alex Abramson

## Abstract

Angiogenesis-promoting macromolecules reduce adverse remodeling and preserve cardiac function in rodents following myocardial infarctions, yet repeatedly fail to translate across length scales in humans. Through mass transport studies in human and swine myocardium, we found that dense, anisotropic myocardial fibers limit therapeutic diffusion and convection to millimeter scales for existing approaches including bolus intramyocardial injections, shear-thinning hydrogels, and epicardial patches. Furthermore, distributions are confined to one dimension along fibers. To increase myocardial drug distribution to centimeter length scales *in vivo* in swine, we engineered a three-dimensional multi-injection drug delivery array. Our device performs up to 40 simultaneous 120 µL injections of functional macromolecules, hydrogels, or mRNA lipid nanoparticles. Injections are precisely placed in relation to fiber alignment, achieving near-complete coverage of the left ventricular myocardium.

## Introduction

Approximately 15% of patients who suffer from a myocardial infarction develop heart failure within one year, primarily due to the formation of scar tissue in the left ventricular myocardium (*1*). Surgical interventions reduce cardiomyocyte necrosis in the myocardium by mechanically restoring blood flow to the infarct, but these interventions still lead to heart failure. Alternatively, macromolecules such as vascular endothelial growth factor (VEGF) aim to reduce scar tissue formation by stimulating angiogenesis (*2-4*). Newly formed blood vessels reduce myocardial scarring in small animal models by increasing nutrient supply to the infarcted area and dampening local inflammation. However, in swine and human clinical trials, VEGF and other therapeutics delivered by intravenous infusions, intracoronary injections, and epicardial patches fail to reduce scar tissue formation or sufficiently improve cardiac function compared to controls (*5-11*). Here, we demonstrate that existing delivery methods for myocardial infarction pharmaceuticals fail to ensure therapeutically relevant concentrations throughout the entire impacted left ventricular myocardium of large animals and humans.

Inherent limitations in diffusion and convection caused by tightly packed myocardial muscle fibers prevent timely transport of VEGF into the infarct, and this delay is exacerbated by size differences between rodent and human hearts. While rodent infarcts are on the millimeter length scale, those in large animals reach centimeter length scales (*12, 13*). Consequently, estimated diffusion timescales of VEGF from patches and depots across millimeter-scale heart walls and infarcts of small animals are approximately 24 hours. Comparatively, over 1 month is required for VEGF to diffuse across centimeter-scale infarcts of large animals (*12, 14*). Myocardial scarring takes place within 1–2 weeks of a myocardial infarction, and VEGF must be delivered within the first few days of a myocardial infarction to be effective (*15-17*). In addition, VEGF is cleared rapidly, as it has a half-life of 30–45 minutes (*18, 19*).

Alternatively, multiple intramyocardial injections can rapidly deliver therapeutic doses of angiogenesis-promoting macromolecules directly into and across the infarcted area, with numerous clinical trials reporting no or few treatment-related serious adverse events (*11, 20-24*). However, without accurate models for predicting injectate distribution in the myocardium, we show that current injection parameters such as volumes, flowrates, and injection site locations, lead to poor distributions by incorrectly assuming isotropic convective and diffusive transport. Using model injections into the left ventricular myocardium of *ex vivo* porcine and diseased human tissue, we found that injected fluids flow predominantly in one direction, along the myocardial fibers. Long, thin cylindrical channels an order of magnitude longer than wide formed due to the highly specific orientations and tight packing of these fibers (*25, 26*). Additionally, manual intramyocardial injections suffer from challenges in accurate needle placement into the myocardium and highly variable flowrates that cause drug leakage from the injection site. We show that current multi-injection procedures result in partial, inefficient drug distribution in the infarct as well as preventable drug leakage out of the myocardium.

Here, we report RISE-LV, The Rapid Injection System for the Entire Left Ventricle. RISE-LV is a multi-needle epicardial injector for rapid and precise macromolecular delivery into the entire left ventricular myocardium during a sternotomy (Fig. 1, A and B). RISE-LV possesses an array of up to 40 needles carefully placed in three-dimensional space to enable macromolecular delivery to the entire left ventricular myocardium without injectate overlap. It simultaneously injects fluid at 60 µL/min into each of the injection sites. This flowrate is slow enough to reduce backflow leakage, while simultaneous injections reduce surgical suite application time to 5-8 minutes. *In vivo* in a porcine model, we show that changing injection parameters away from those currently used in clinical trials can significantly increase tissue coverage. Our RISE-LV multi-needle injection array delivered macromolecules to an order of magnitude greater tissue volume than existing delivery methods such as bolus injections and epicardial patches, and it enables near-complete coverage of the entire left ventricular myocardium (table S1-3). We show that RISE-LV is compatible with aqueous formulations of functional macromolecules, mRNA-encapsulating lipid nanoparticles, and shear thinning hydrogels (*27*), enabling improved tissue coverage for formulations designed to provide long-lasting effects.

**Fig. 1.**
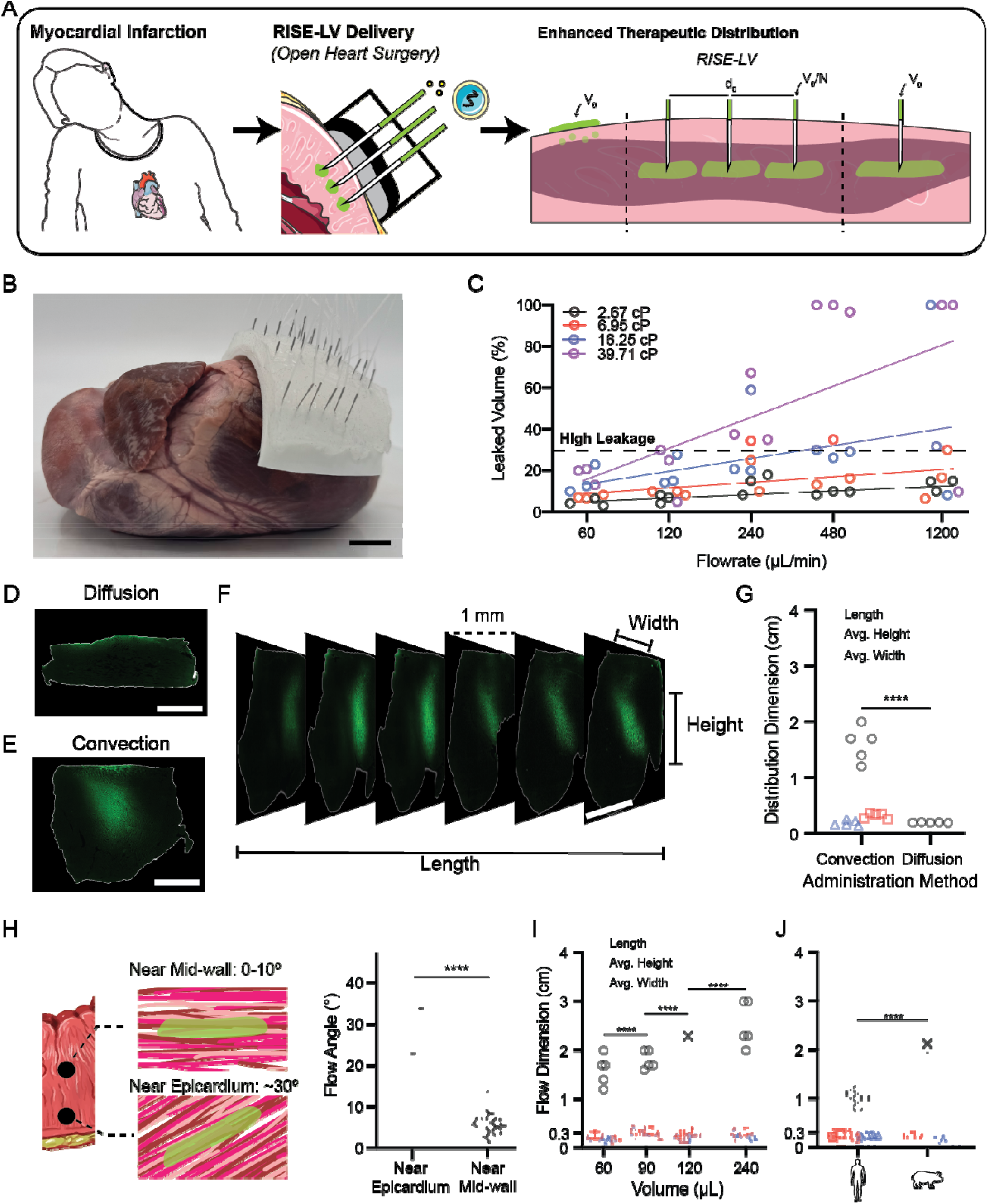
Multi-injection arrays overcome limited diffusive and convective macromolecular transport. (**A**) RISE-LV deployment during open heart surgery. (**B**) RISE-LV prototype. (**C**) Leakage characterization of solutions with varying viscosities and flowrates. *n* = 3 injections. (**D-E**) Representative fluorescent microscopy images showing distribution of 40 kDa FITC-dextran by diffusion (**D**) and convection (**E**). Bright green represents FITC-dextran and tissue boundaries are outlined in white. (**F**) Representative stack of fluorescent microscopy images following a FITC-dextran injection. (**G**) Distribution dimensions of FITC-dextran from different administration methods. *n* = 5 tissue samples. Statistical analysis by a one-way ANOVA followed by Tukey’s multiple comparisons test. \*\*\*\**P* < 0.0001. (**H**) Myocardial fiber orientation angles near the mid-wall and epicardium, and flow angles following injection into each layer. *n* = 7-17 injections. (**I**) Injectate distribution in the myocardium at different injection volumes. *n* = 5 injections. Statistical analysis by a one-way ANOVA followed by Tukey’s multiple comparisons test. \*\*\*\**P* < 0.0001. (**J**) Flow dimensions in *ex vivo* human and porcine myocardium. *n* = 5-12 injections. Statistical analysis by an unpaired two-tailed Student’s *t* test. \*\*\*\**P* < 0.0001. Scale bars = 2.5 cm (B), 2 mm (D, E), and 1 cm (F).

## Results

### Characterizing Macromolecular Transport in the Myocardium

We characterized macromolecular diffusion and convection in both *ex vivo* healthy porcine and diseased human left ventricular myocardium to determine injection parameters and sites that enable complete impacted tissue coverage following a myocardial infarction.

Injectate viscosity and flowrate directly contributed to the proportion of drug that remained in the tissue following an injection. We delivered fluids with a range of viscosities comparable to therapeutic modalities such as protein solutions and injectable hydrogels into porcine myocardium. At all viscosities, low 60-120 µL/min flowrates attainable only by injection pumps resulted in 2-5x lower leakage volumes than the 240-1200 µL/min manual injection flowrates typically performed during clinical trials (Fig. 1C) (*28*). Additional experiments therefore only utilized flowrates between 60-120 µL/min to minimize leakage.

Due to tight muscle fiber packing, diffusion from epicardial patches or from bolus injection sites yielded minimal mass transport over a 24-hour period in *ex vivo* porcine left ventricular myocardium. We utilized 40 kDa FITC-dextran as a fluorescent, visualizable proxy of the 38.2 kDa VEGF_165_ homodimer, due to similarities in molecular weight. Histology and fluorescent microscopy enabled visualization of molecular distribution within the myocardium (Fig. 1 D to G). After 24 hours, FITC-dextran diffused only 0.19 ± 0.04 cm, just 11.5% of the mean left ventricular free wall thickness in diseased humans (Fig. 1G) (*29*). This distance is in line with estimates based on the standard diffusion timescale estimation formula *L*^*2*^*/D*, where *L* denotes diffusion length and *D* denotes the diffusion coefficient. Here, we estimate *D* to be on the order of 10^-10^ m^2^/s for our chosen molecular weight (*4*). This formula predicts timescales of > 4 weeks for diffusion across the entire left ventricular free wall. On the other hand, intramyocardial injections of the same volume transported FITC-dextran 1-2 cm within 5 minutes (Fig. 1 E to G and fig. S1). Following the injection, macromolecules diffused just 0.2 ± 0.05 cm (mean ± s.d.) in each direction from the injection site over a 24-hour period (Fig. 1G).

Injected fluids formed long, narrow channels in the myocardium. We hypothesized that they formed this shape rather than spherical boluses due to the tight packing and highly ordered, anisotropic arrangement of myocardial fibers. When we injected FITC-dextran solutions into different depths of the myocardium, where fibers are predictably oriented at different angles, we found that the flow angles matched local fiber angles (Fig. 1H). These results align with prior research showing 10x greater permeability along myocardial fibers than in other directions (*25, 26*). Due to slow diffusion, anisotropic injectate distributions remain even after 24 hours in the tissue. Because single injections primarily flow in in one dimension, it is impossible to cover the entire left ventricular myocardium without performing multiple injections extending perpendicular to the fiber alignment.

The tightly packed myocardial fibers enabled but constrained convective flow parallel to their alignment. Injecting larger volumes affected only flow length and had no impact on height or width distributions (Fig. 1I), but gains were lower than the expected linear relationship. This is consistent with observations of injections in other types of tissue such as the brain (*30*). Changing the flowrate and needle insertion angle resulted in no statistically significant changes to the flow dimensions (fig. S2). Due to the limited ability to distribute injectate along the fiber lengths, multiple injections of small volumes extending parallel to the fibers are required to cover the entire myocardium.

Using diseased human left ventricular myocardium samples from five subjects with cardiomyopathies (table S4), we noted that existing cardiomyopathies may further reduce injectate distribution in the myocardium. In all samples we observed the same anisotropic injectate distribution (Fig. 1J), despite greater collagen buildup compared to porcine tissue (fig. S3). This is in line with previous studies showing that newly deposited collagen fibers are oriented at the same angles as adjacent myocardial fibers (*31*). Injections into diseased human tissue demonstrated 40% decreased injectate lengths compared to *ex vivo* porcine tissue.

By precisely aligning an array of needles to the myocardial fibers, multi-point delivery of small volumes can significantly improve tissue distributions compared to bolus injections. Utilizing microcomputed tomography (micro-CT), we analyzed the distribution of phosphotungstic acid, a macromolecular contrast agent, mixed with an injectable hydrogel developed for intracardiac injections by Cohen et al. (*27*). We compared contrast agent distribution from an epicardial patch, a single injection of 540 µL, and nine 60 µL injections in a 3 x 3 array (Fig. 2, A to C). While a single injection elicited a 4x greater volume of distribution compared to an epicardial patch, multiple injections with the same total volume further increased the volume of distribution 3x compared to bolus delivery (Fig. 2D).

**Fig. 2.**
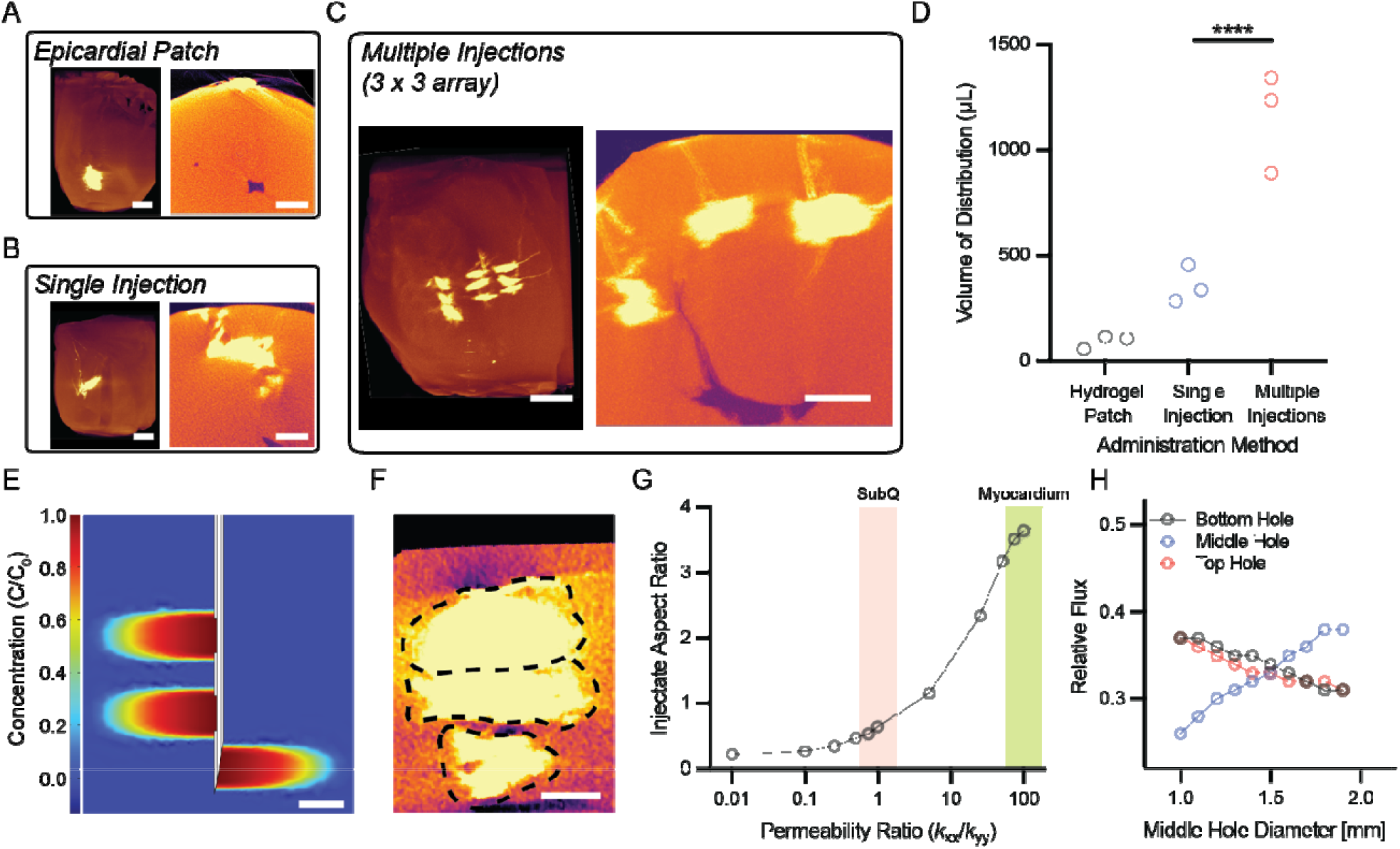
Multi-Point injections aligned with myocardial fibers improve injectate distributions. (**A, B, C**) Longitudinal (left) and axial (right) micro-CT view of the heart following delivery of a shear-thinning, extended-release hydrogel loaded with phosphotungstic acid using either an epicardial patch (**A**), a single injection of 540 µL (**B**), and nine injections of 60 µL in a 3 x 3 array (**C**). (**D**) Volume of distribution from micro-CT. *n* = 3 hearts. Statistical analysis by a one-way ANOVA followed by Tukey’s multiple comparisons test. \*\*\*\**P* < 0.0001. (**E**) A COMSOL simulation of injection into a myocardium demonstrating equal flux out of all 3 holes of a fenestrated needle. (**F**) Micro-CT scan of the myocardium following phosphotungstic acid injection with a fenestrated needle. (**G**) The 2D permeability tensor affects aspect ratios of injectate distribution in *ex vivo* tissue models and in COMSOL simulations. (**H**) Hole size optimization in COMSOL achieves equal relative flux from all 3 holes of a fenestrated needle. Scale bars = 1 cm (A, B, C) and 2 mm (E, F).

While individual injectate height dimensions from 120 µL injections are only ∼ 20% of the total myocardium thickness, fenestrated needles can be used to increase total injectate height distribution (Fig. 2, E and F). Prior to determining sizes and locations of new holes, we built a 2D COMSOL model that exhibited similar injectate distributions to those obtained from *ex vivo* experiments. This was achieved by altering components of the permeability tensor to reflect the anisotropicity of myocardial tissue (Fig. 2G). We performed simulations to obtain hole size parameters that allowed equal flowrates out of all needle holes into the myocardium (Fig. 2H). To demonstrate the effects of multi-hole injections, we etched two new holes into 27G hypodermic needles using a laser. By injecting phosphotungstic acid into the myocardium with these needles and visualizing the distribution with micro-CT, we show that individual injectate height can be predictably increased using this method without increasing the number of injected needles.

### Device for Simultaneous Injections into Beating Hearts

RISE-LV is an injection system that can be used during open heart surgery to deliver macromolecules to the entire left ventricular myocardium (Fig. 3, A to C). It is comprised of up to 40 injection needles depending on the size of the heart. Its hard backing places each needle in a location relative to the myocardial fiber alignment that minimizes injectate overlap while maximizing tissue coverage. A backing handle provides a surface to push the device onto the heart. The device possesses a high-friction, elastomeric layer to prevent the needles from tissue ejection during insertion into the heart. Furthermore, it employs a strap that secures the device and needles in place while performing the injections into a beating heart.

**Fig. 3.**
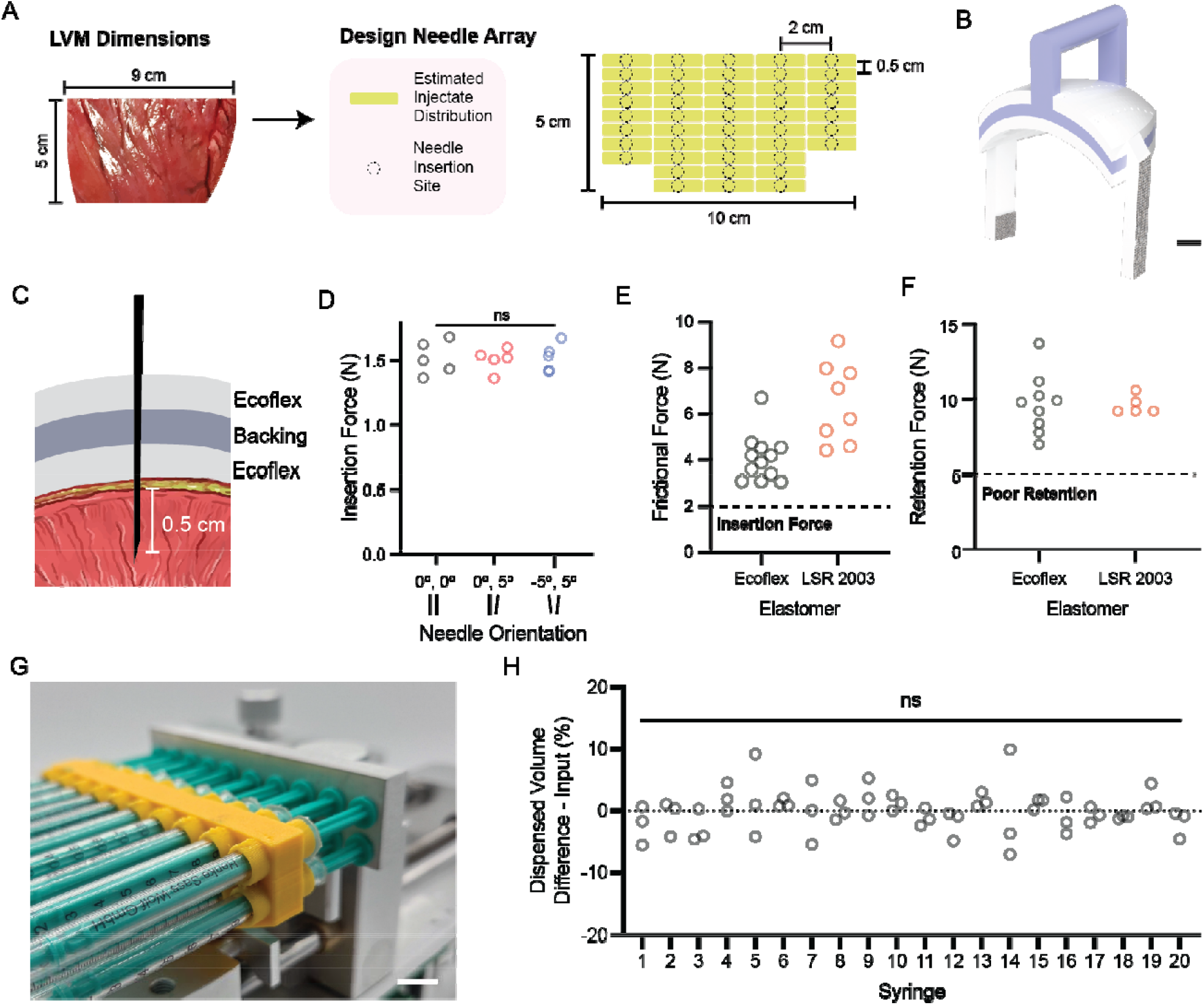
Engineering an intramyocardial multi-injection array device. (**A**) Needle array dimensions are defined by left ventricular myocardium (LVM) dimensions and *ex vivo* injectate distribution studies. (**B**) Computer-aided design model of RISE-LV. White, purple, and grey represent Ecoflex, the hard backing, and Velcro, respectively. (**C**) A cross-section of a needle in RISE-LV. (**D**) Minimum insertion force required to insert 2 needles simultaneously into the myocardium does not change at different needle orientations. *n* = 5 tissue insertion tests. Statistical analysis by a one-way ANOVA. (**E**) Minimum frictional force experienced by a single needle embedded in Ecoflex or LSR 2003 overcomes insertion forces into the myocardium. *n* = 8-12 needles. (**F**) Minimum retention force provided by Ecoflex or LSR 2003 straps is greater than ejection forces imposed from heart beats. *n* = 5-8 devices. (**G**) Multi-injection setup. The syringe rack (yellow) holds up to 20 needles in a 2 x 10 array. All plungers are simultaneously pushed with an aluminum slab. (**H**) Discrepancy between dispensed volume and programmed dispensed volume from 20 syringes. *n* = 3 injections. Statistical analysis by a one-way ANOVA.

The RISE-LV array allows simultaneous needle insertion into the mid-wall of the myocardium in a pattern that prevents overlapping injectates. Based on *ex vivo* flow dimensions, RISE-LV places needles 1.5-2 cm apart horizontally and 0.5 cm apart vertically. Each needle protrudes 5 mm, such that all needles are inserted into the mid-wall. RISE-LV delivers to the mid-wall to reduce the risk of perforation of the left ventricular free wall as well as avoid insufficient needle penetration. This allows the device to function as intended even when accounting for up to 25% variabilities in myocardial thicknesses amongst individuals (*29*).

A handle allows the user to easily apply sufficient force to insert all needles. To minimize insertion force, all needles must be oriented vertically, along the axis that RISE-LV is pushed onto the heart. The 5 mm thickness of the backing and 0.7 mm diameter of needle holes limit the vertical tilt angles of needles to < 5º (fig. S4). We found no significant difference between forces required to insert vertical needles and those tilted at 5º (fig. S5). Similar results were found when simultaneously inserting 2 needles (Fig. 3D). A force of 24.32 ± 2.55 N is required to insert 40 needles simultaneously into *ex vivo* porcine hearts (mean ± s.d., n = 5 devices and hearts), similar to the force required to push a light shopping cart.

By embedding all needles in an elastomer, RISE-LV prevents needle displacement during insertion into the myocardium. We used a minimum frictional force benchmark of 2 N, based on our insertion force results as well as previous studies (*32*). To measure the friction provided by elastomer layers, we used a mechanical test stand to pull on needles vertically at 1 cm/s, the approximate speed at which the heart contracts and expands. Needles inserted into pre-cured elastomers with thicknesses up to 2 cm failed to achieve retention forces greater than 2 N (fig. S6). Alternatively, when elastomers were directly cured onto the needles, a minimum force 2x greater than the benchmark was required to remove each needle (Fig. 3E). We used Ecoflex (Ecoflex 00-30) for prototyping and also showed that a USP Class VI certified biocompatible alternative, LSR 2003, provided even greater retention forces without altering the fabrication process.

Elastomeric straps with Velcro secure RISE-LV onto the epicardium of a beating heart and prevent needle ejection during heartbeats (Fig. 3F). While there is no FDA-mandated standard for minimum retention force of cardiac implanted devices, we sought to achieve retention forces greater than 5 N. This traction force requirement is mandated by the EN 45502-2-1 and ISO 14708-1 standards for the mechanical integrity of active implantable cardiac pacemaker leads. Elastomeric straps provide rapid applicability and reversibility, minimal invasiveness, and biocompatibility; these criteria rule out conventional methods such as adhesives, suturing, and suction. An Ecoflex elastomeric strap on one side of the backing was stretched around the heart until a strain of 0.5 was reached and then attached to Velcro on the opposite side. At this strain, the theoretical maximum pressure applied onto the heart by the strap was 300 Pa. This value is 25% lower than pressures previously evaluated to be safe when applied to the heart over months in clinical trials (*17, 18*). When the strap was wrapped around the heart, the minimum force required to lift RISE-LV 0.5 cm, enough to displace needles from the myocardium, was 9.70 ± 1.88 N (Fig. 3F). This force satisfies the traction force requirements and is also greater than the force required to remove FDA approved surgical suction tools such as the Medtronic Starfish from the heart (*16*). Similar values were achieved with LSR 2003, when two straps were stretched around the heart until a strain of 0.05 was reached (Fig. 3F). LSR 2003 exerts the same pressure as Ecoflex at this strain, due to its greater Young’s modulus.

Manually performing up to 40 precisely placed intramyocardial injections of 120 µL at 60 µL/min is both physically infeasible and exceptionally time-consuming. We developed a custom syringe rack that holds 20 syringes per bay (Fig. 3, G and H, and fig. S7), enabling up to 40 simultaneous injections with a 2-bay syringe pump. A 1 cm thick aluminum slab cut into the same shape as the rack enables simultaneous plunger displacement and fluid dispensation from all syringes of each bay. The mean dispensed volume of fluid from each of the 20 syringes was within 0.1% error of the specified volume, with no statistically significant differences amongst the means of syringes at each rack position (Fig. 3H).

### *In vivo* Deployment of RISE-LV in Swine

*In vivo* in swine, we deployed RISE-LV following a sternotomy (Fig. 4, A and B). By enabling simultaneous delivery, RISE-LV allowed the surgeon to complete all injections within a 5-8 minute timeframe. The elastomer layers of RISE-LV provided sufficient retention force to prevent needle displacement and ejection during the surgery. Elastomeric straps that wrapped RISE-LV around the beating heart provided additional retention, but the device also remained on the heart using only the applied force by the surgeon.

**Fig. 4.**
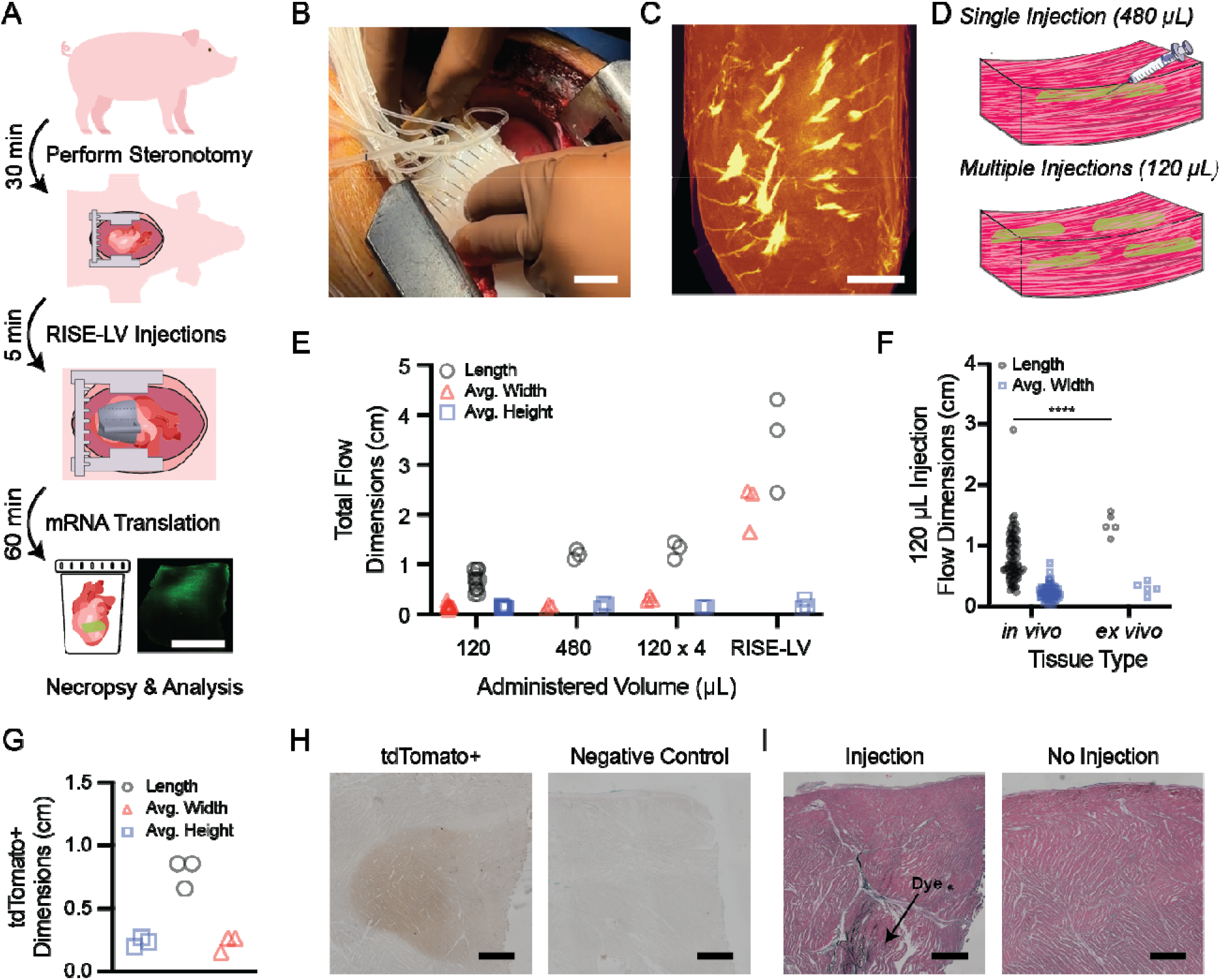
*In vivo* deployment of RISE-LV in swine. (**A**) Timeline of swine experiments. (**B**) RISE-LV is delivered onto a heart following a sternotomy. (**C**) Micro-CT scan of the left ventricular free wall following phosphotungstic acid injection with RISE-LV. (**D**) Single, bolus injections of 480 µL were compared to four 120 µL injections in a 2 x 2 array. (**E**) Comparison of total flow dimensions from different total administered volumes. *n* = 3-12 injections. (**F**) Flow dimensions of phosphotungstic acid channels in the myocardium in *in vivo* and *ex vivo* porcine myocardium following 120 µL injections. *n* = 5-87. Statistical analysis by an unpaired two-tailed Student’s *t* test. \*\*\*\**P* < 0.0001. (**G**) Following delivery of mRNA lipid nanoparticles encoding for tdTomato using RISE-LV the translated protein is distributed anisotropically in the myocardium. (**H**) Representative immunohistochemistry stain against tdTomato. Brown represents tdTomato. No tdTomato expression was detected in the negative control group. (**I**) Representative Hematoxylin & Eosin stain of the myocardium following RISE-LV administration compared to a negative control. We found no visually assessable damage in either needle insertion sites or negative control groups. Scale bars = 2 cm (B), 1 cm (C), and 1 mm (G-H).

Our experiments demonstrated the need to perform 26-27 injections across the left ventricle of each animal’s heart to enable macromolecule access over the entire tissue volume. Injection numbers varied based on heart size, and the swine used for the *in vivo* experiment had smaller hearts than the ones used for *ex vivo* studies. Utilizing micro-CT, we visualized RISE-LV delivery of aqueous phosphotungstic acid solution (Fig. 4C and fig. S8). Consistent with results from *ex vivo* porcine tissue, we observed narrow channels forming along myocardial fibers. By implementing the array rather than performing bolus injections, the narrow injectate distributions enabled near-complete coverage of the left ventricular myocardium. By splitting up our drug volume into a multi-injection array, RISE-LV distributed injectate to a 2.6x greater tissue volume than a single injection of equal total volume (Fig. 4 D, E, and fig. S9). The complete RISE-LV array provided 10x the tissue coverage compared to a bolus 480 µL injection. The total injectate volume administered in our array was 3.16 ± 0.06 mL (mean ± s.d.) over 26-27 injections of 120 µL, spaced 1.5 cm apart parallel to myocardial fibers, and spaced 0.5 cm apart perpendicular to myocardial fibers. Previous clinical trials delivered 1-6 mL of total volume via arbitrarily placed 100-1000 µL injections (table S3) (*33-35*). Our study demonstrates the need to perform precisely placed injections to avoid tissue overlap while simultaneously performing large numbers of low volume injections to increase the total tissue coverage. Compared to *ex vivo* measurements, this need is even more exacerbated *in vivo*, where there was a 35% decrease in flow length (Fig. 4F).

We also delivered functional mRNA-encapsulating lipid nanoparticles using RISE-LV. tdTomato encoding mRNA demonstrated protein translation in cardiomyocytes 1 hour following administration. Immunohistochemistry stains showed that the translated protein was anisotropically distributed in the tissue, akin to the dye distributions from our above *in vivo* experiments (Fig. 4, G, H, and fig. S10). Therefore, mRNA therapeutic distribution also benefits from the precise injection arrays enabled by RISE-LV administration.

While all animals were euthanized within 1 hour following the injections, no animal displayed signs of severe trauma that required immediate euthanasia during or immediately after the injections. Hematoxylin and Eosin stains also show no apparent damage near needle insertion sites (Fig. 4I). This safety profile is in line with prior large animal studies and clinical trials where pharmaceuticals were injected directly into the heart (*11, 20, 21*).

## Discussion

We report the development of RISE-LV, an epicardial multi-needle injector that rapidly delivers macromolecules by convection to the entire left ventricular myocardium in just 5-8 minutes during open heart surgery. We show that drug delivery methods typically used pre-clinically or in clinical trials such as epicardial patches, single bolus injections, or randomly placed injections deliver macromolecules to 62-92% less tissue compared to RISE-LV. Therapeutics can only function when they reach their target site, and we demonstrate that tissue-scale transport is a principal bottleneck in cardiac drug delivery. RISE-LV is a scalable solution designed specifically to solve this problem, enabling near-complete tissue coverage *in vivo* in swine left ventricular myocardium.

Our device utilizes optimized injection parameters and the anisotropic distribution of injectates along myocardial fibers to engineer a 26 to 40-point array that enables this improved tissue distribution *in vivo* in a porcine model. We observed that injectates form narrow channels along myocardial fibers in all trials. The length dimension, however, changed ∼40% when comparing results between human and swine as well as between *ex vivo* and *in vivo* tissues. This could be attributed to a combination of variables such as tissue hydration states, tissue interstitial pressures, heart motions, and permeability differences due to freeze-thaw cycles. Human trials will be required to understand the potential altered mass transport distributions in live human myocardial tissue compared to our *in vivo* swine studies. These human trials should also assess the long-term safety and efficacy of multi-needle array injections, as our study only looked at the acute impacts of RISE-LV administration.

We demonstrate the delivery of multiple injectates, including functional mRNA and shear-thinning hydrogels, using RISE-LV. Further studies will be required to understand the impact of improved drug distribution on therapeutic efficacy, both in large animal and human trials. Injectate distribution may have varying impacts on therapies depending on their method of action. Currently researchers are studying peptides, proteins, nucleic acids, cell therapies, and injectable hydrogels for cardiac tissue regeneration. All of these modalities are compatible with RISE-LV.

While no animal experienced severe trauma related to the RISE-LV procedure, one limitation of the device is the invasive surgery required to perform device delivery. Reducing invasiveness, such as performing transendocardial injections through catheters, may allow for more patients to benefit from this technology. Currently, ∼10% of myocardial infarction patients undergo a coronary artery bypass graft surgery that is compatible with RISE-LV administration (*1*). However, while injectate distributions in the myocardium remain the same regardless of the needle entry location, catheter delivery requires sequential injections that increase administration times and add uncertainty to injection placements. Overall, our study demonstrates the importance of injection location, volume, and flowrate when administering drugs to the myocardium and uses this information to engineer a technology that significantly improves drug distribution compared to the current pre-clinical and clinical standards.

## Supporting information

All data from thsi study.

Python code used for data analysis.

## Acknowledgements

We thank members of the Georgia Tech Parker H. Petit Institute for Bioengineering and Bioscience Core Facilities for the discussion and technical support: L. Krishnan (Biomechanics and Microcomputed Tomography Cores), S. Hsieh (Optical Microscopy Core), A. Asberry (Histology Core), and D.J. Alexander (Histology Core); and members of the Emory University Department of Animal Resources for their help during the swine surgeries. We thank R. Shad for his insight into myocardial infarction treatments.

## Author contributions

J.P. and A.A. designed all the experiments. J.P., A.R., E.B., A.M., H.M., Y.A. performed the *ex vivo* porcine tissue experiments. J.P., E.B., A.M., Y.A., and H.M. performed COMSOL simulations. J.P. performed the *ex vivo* human tissue experiments. R.G., and A.M.A. fabricated and characterized the mRNA-encapsulating lipid nanoparticles. J.E.D. supervised the lipid nanoparticle work. M.B. and J.L.C. performed the swine surgeries. J.J., S.H., J.Z.D., and M.C. assisted the swine experiments. J.P., E.B., A.M., and Y.A. performed analyses of the porcine tissue. A.A. provided the funding and supervised the project. All authors contributed to the writing of the manuscript.

## Data and materials availability

All data associated with this study are present in the paper or the Supplementary Materials.

## Code availability

The code developed for this study is available for download from the supplementary materials section of the paper.

## Supplementary Materials

Materials and Methods

Figs. S1 to S15

Table S1 to S4

Supplementary

References 36-37

Excel Data File

Python Code

