## Supplementary material for "Multi-point convective delivery overcomes mass transport barriers for myocardial therapeutics": Python code used for data analysis.: Avg. Width & Height.py

```
# 1. Import processing modules
import cv2 as cv
import numpy as np
from matplotlib import pyplot as plt
import os
import pandas as pd
import re

# Copyrighted by Jihoon Park. Code written with the assistance of Gemini.
# This python script evaluates the maximum width, average width, maximum height, and average height of all individual thresholded images in pixels. These values are stored in a spreadsheet and can then be converted to millimeters with their original scale bars.

# Conditional import for Google Colab compatibility
try:
    from google.colab.patches import cv2_imshow
except ImportError:
    def cv2_imshow(img):
        # Note: In local environments, cv.imshow requires a loop or waitKey to stay open
        cv.imshow("Image", img)
        cv.waitKey(1) 
        cv.destroyAllWindows()

# Specify folder path containing thresholded images. Compatible file extensions are .jpg, .png, and .jpeg
# 2. Define paths
folder_path = 
output_folder = os.path.join(folder_path, "plots")
excel_output_folder = os.path.dirname(os.path.dirname(folder_path))

# Ensure output folders exist
os.makedirs(output_folder, exist_ok=True)
os.makedirs(excel_output_folder, exist_ok=True)

# Get a list of all image files in the folder
image_files = [f for f in os.listdir(folder_path) if f.lower().endswith((".jpg", ".jpeg", ".png"))]

# Create an empty DataFrame to store the data
columns = [
    "Position (mm)", "Width (Px)", "Avg. Height (Px)", "Max. Height (Px)",
    "Height (Px)", "Avg. Width (Px)", "Max. Width (Px)", "Scale Bar", "Avg. Height (mm)", "Avg. Width (mm)"
]
data_list = []

# This block calculates the max width, average width, max height, and average height of all black pixels in a thresholded image.
# 3. Process each image
for image_file in image_files:
    image_path = os.path.join(folder_path, image_file)
    print(f"Processing: {image_file}")

    input_image = cv.imread(image_path, cv.IMREAD_GRAYSCALE)

    if input_image is None:
        print(f"Error: Unable to load {image_file}")
        continue

    # Copy and rotate the image
    img = input_image.copy()
    rot_img = cv.rotate(img, cv.ROTATE_90_COUNTERCLOCKWISE)

    # Thresholding (Using user logic: Binary Inv catches objects on black background)
    _, thresh1 = cv.threshold(img, 1, 255, cv.THRESH_BINARY_INV)
    _, thresh2 = cv.threshold(rot_img, 1, 255, cv.THRESH_BINARY_INV)

    # Find contours
    contours, _ = cv.findContours(thresh1, cv.RETR_EXTERNAL, cv.CHAIN_APPROX_SIMPLE)
    rot_contours, _ = cv.findContours(thresh2, cv.RETR_EXTERNAL, cv.CHAIN_APPROX_SIMPLE)

    if not contours or not rot_contours:
        print(f"No contours found in {image_file}")
        continue

    # Create masks and draw contours
    h, w = thresh1.shape
    result_mask = np.zeros((h, w), np.uint8)
    cv.drawContours(result_mask, contours, -1, 255, -1)

    rot_h, rot_w = thresh2.shape
    rot_result_mask = np.zeros((rot_h, rot_w), np.uint8)
    cv.drawContours(rot_result_mask, rot_contours, -1, 255, -1)

    # Calculate depth and width metrics
    depth = [cv.countNonZero(result_mask[:, j]) for j in range(w)]
    width = [cv.countNonZero(rot_result_mask[:, j]) for j in range(rot_w)]

    if not any(depth) or not any(width):
        print(f"Skipping {image_file} due to empty depth/width arrays")
        continue

    length_x = sum(1 for d in depth if d > 0)
    average_h = sum(depth) / length_x if length_x > 0 else 0
    maximum_h = max(depth)

    width_y = sum(1 for w in width if w > 0)
    average_h2 = sum(width) / width_y if width_y > 0 else 0
    maximum_h2 = max(width)

    # Store data in a list (Faster than pd.concat)
    data_list.append({
        "Position (mm)": image_file, # Keep full filename for now
        "Width (Px)": length_x,
        "Avg. Height (Px)": average_h,
        "Max. Height (Px)": maximum_h,
        "Height (Px)": width_y,
        "Avg. Width (Px)": average_h2,
        "Max. Width (Px)": maximum_h2
    })

# Height and width at all x and y values (horizontal and vertical axes) are plotted; verify accuracy of code with these plots.
    # Plotting
    plt.figure(figsize=(6, 4))
    plt.plot(range(len(depth)), depth, color="red", lw=1)
    plt.title(f"Height vs Width: {image_file}")
    plt.xlabel("Flow Width, x")
    plt.ylabel("Flow Height, h")
    plot1_path = os.path.join(output_folder, f"{os.path.splitext(image_file)[0]}_plot1.png")
    plt.savefig(plot1_path)
    plt.close()

    plt.figure(figsize=(6, 4))
    plt.plot(range(len(width)), width, color="blue", lw=1)
    plt.title(f"Width vs Height: {image_file}")
    plt.xlabel("Flow Height, y")
    plt.ylabel("Flow Width, h2")
    plot2_path = os.path.join(output_folder, f"{os.path.splitext(image_file)[0]}_plot2.png")
    plt.savefig(plot2_path)
    plt.close()

# 4. Final Data Organization and Hierarchical Sorting
data = pd.DataFrame(data_list)

def get_sort_key(filename):
    """Finds all numbers in the filename to sort by (e.g., [1, 4, 0])."""
    numbers = re.findall(r"-?\d+\.?\d*", filename)
    return [float(n) for n in numbers]

def clean_name(filename):
    """Removes extension for a cleaner Excel sheet."""
    return os.path.splitext(filename)[0]

if not data.empty:
    # Create the temporary sorting column
    data['Sort_Key'] = data['Position (mm)'].apply(get_sort_key)
    
    # Sort by the numerical list [FirstNum, SecondNum, ThirdNum]
    data = data.sort_values(by='Sort_Key').reset_index(drop=True)
    
    # Clean the display name AFTER sorting
    data['Position (mm)'] = data['Position (mm)'].apply(clean_name)
    
    # Remove temporary column
    data = data.drop(columns=['Sort_Key'])

    # Save data to Excel
    excel_file_path = os.path.join(excel_output_folder, "image_data.xlsx")
    data.to_excel(excel_file_path, index=False)
    print(f"Success! Data saved to {excel_file_path}")
else:
    print("No data collected. Excel file not created.")
```
