## Supplementary material for "Multi-point convective delivery overcomes mass transport barriers for myocardial therapeutics": Python code used for data analysis.: Volume of Distribution.py

```
import os
import cv2
import numpy as np
import pyvista as pv
from scipy.ndimage import distance_transform_edt

# Copyrighted by Jihoon Park. Code written with the assistance of Gemini.
# This python script places (2D xy) thresholded images along the z axis, interpolates between the images, and calculates the volume of distribution.

# --- 1. Settings ---
# Specify path with thresholded images here; compatible file extensions are .jpg, .png, and .jpeg
folder_path = 

# Specify the first and last z coordinates of the image stack. e.g., the first image here is placed at -7.5 mm, and the last image is placed at 7.5 mm.
# Physical boundaries in mm (100 px/mm scale)
Z_START_PX = -750
Z_END_PX = 750
Z_SPAN_PX = Z_END_PX - Z_START_PX 

PINK_OPACITY = 0.2   
TISSUE_OPACITY = 1.0  
SLICES_PER_UNIT = 10  # Density of interpolation

# --- Helper Function: Signed Distance Field ---
def get_sdf(img, threshold=254):
    mask = img < threshold
    if not np.any(mask): return np.ones_like(img, dtype=float) * 1000
    if np.all(mask): return np.ones_like(img, dtype=float) * -1000
    dist_in = distance_transform_edt(mask)
    dist_out = distance_transform_edt(~mask)
    return dist_out - dist_in

# --- 2. Load and Parse Images ---
valid_exts = ('.jpg', '.png', '.jpeg')
image_files = sorted([f for f in os.listdir(folder_path) if f.lower().endswith(valid_exts)])

image_data = []
for f in image_files:
    try:
        z_mm = float(os.path.splitext(f)[0])
        img = cv2.imread(os.path.join(folder_path, f), cv2.IMREAD_GRAYSCALE)
        if img is not None:
            image_data.append({'z': z_mm, 'sdf': get_sdf(img)})
    except ValueError:
        continue

image_data.sort(key=lambda x: x['z'])
target_h, target_w = image_data[0]['sdf'].shape
total_slices = int(15 * SLICES_PER_UNIT) + 1
known_zs = [d['z'] for d in image_data]

# This block smooths the reconstructed image.

# --- 3. Build Smooth Volume ---
volume_sdf = np.zeros((target_h, target_w, total_slices), dtype=float)

for k in range(total_slices):
    current_z_mm = -7.5 + (k / SLICES_PER_UNIT)
    
    match_idx = -1
    for i, z_val in enumerate(known_zs):
        if np.isclose(current_z_mm, z_val, atol=1e-5):
            match_idx = i
            break
            
    if match_idx != -1:
        volume_sdf[:, :, k] = image_data[match_idx]['sdf']
    elif current_z_mm < known_zs[0] or current_z_mm > known_zs[-1]:
        volume_sdf[:, :, k] = 100.0 
    else:
        idx = np.searchsorted(known_zs, current_z_mm)
        z_a, z_b = known_zs[idx-1], known_zs[idx]
        sdf_a, sdf_b = image_data[idx-1]['sdf'], image_data[idx]['sdf']
        weight_b = (current_z_mm - z_a) / (z_b - z_a)
        volume_sdf[:, :, k] = (1.0 - weight_b) * sdf_a + weight_b * sdf_b

# This block creates the grid in which images are stacked.

# --- 4. Centering and Grid Setup ---
volume_xyz = np.transpose(volume_sdf, (1, 0, 2))
grid = pv.ImageData(dimensions=volume_xyz.shape)
grid.origin = (-target_w / 2, -target_h / 2, Z_START_PX)
grid.spacing = (1, 1, Z_SPAN_PX / (volume_xyz.shape[2] - 1))
grid.point_data["values"] = volume_xyz.flatten(order="F")

# This block calculates the volume of distribution of the reconstructed stack.

# --- 5. Rendering & Volume Calculation ---
plotter = pv.Plotter(off_screen=True)
plotter.enable_parallel_projection()

# A. The Tissue
tissue_mesh = grid.contour(isosurfaces=[0], scalars="values")
if tissue_mesh.n_points > 0:
    smoothed = tissue_mesh.smooth_taubin(n_iter=30, pass_band=0.05)
    plotter.add_mesh(smoothed, color='yellowgreen', smooth_shading=True, opacity=TISSUE_OPACITY)

    # Volume calculation
    raw_volume_px3 = smoothed.volume
    volume_mm3 = raw_volume_px3 / 1_000_000
    print("-" * 30)
    print(f"CALCULATED VOLUME: {volume_mm3:.6f} mm³")
    print("-" * 30)

# B. Pink Background
empty_voxels = grid.threshold([0.001, 1000], scalars="values")
if empty_voxels.n_points > 0:
    plotter.add_mesh(empty_voxels.extract_surface(), color='pink', opacity=PINK_OPACITY, lighting=False)

# C. Outline First and Last Slices Only
if len(known_zs) >= 2:
    for z_val in [known_zs[0], known_zs[-1]]:
        z_world_px = z_val * 100
        rect = np.array([
            [-target_w/2, -target_h/2, z_world_px],
            [ target_w/2, -target_h/2, z_world_px],
            [ target_w/2,  target_h/2, z_world_px],
            [-target_w/2,  target_h/2, z_world_px],
            [-target_w/2, -target_h/2, z_world_px]
        ])
        plotter.add_mesh(pv.MultipleLines(points=rect), color='red', line_width=4)

# D. Outer Bounding Box
plotter.add_mesh(grid.outline(), color="darkgray", line_width=1.5)
plotter.view_isometric()
plotter.screenshot(os.path.join(folder_path, "3D_Final_Reconstruction.png"), transparent_background=True)
plotter.close()

print(f"Success! Reconstruction complete.")
```
