## Supplementary materials. for "Multi-point convective delivery overcomes mass transport barriers for myocardial therapeutics"

**The PDF file includes:**

Materials and Methods
Figs. S1 to S15
Tables S1 to S4

Table of Contents

**Materials and Methods**

**Supplementary Figures**

*Fig. S1. Injectate distribution after 0 and 24 hours.*

*Fig. S2. Effects of injection flowrate and needle angle on myocardial injectate distribution.*

*Fig. S3. Histological stains of human and porcine myocardium samples.*

*Fig. S4. Design of the backing limiting vertical tilt angle deviation.*

*Fig. S5. Single needle insertion force at 0º and 5º.*

*Fig. S6. Single needle frictional force.*

*Fig. S7. Syringe Rack Design & Characterization.*

*Fig. S8. Micro-CT scans of the left ventricular myocardium of swine dosed in vivo with RISE-LV.*

*Fig. S9. Four + one injection array RISE-LV used in vivo in some studies.*

*Fig. S10. Representative Immunohistochemistry stains against tdTomato. Brown indicates tdTomato.*

*Fig. S11. Setup of diffusion studies.*

*Fig. S12. Injectate visualization by histology and microscopy.*

*Fig. S13. Flow angle calculation by histology and microscopy.*

*Fig. S14. Procedure used to assemble RISE-LV prototypes.*

*Fig. S15. Formulation components and quality control of formulated tdTomato mRNA-encapsulating lipid nanoparticles.*

**Supplementary Tables**

*Table S1. Summary of mass transport dimensions in porcine hearts.*

*Table S2. Preclinical device comparisons for cardiac drug delivery.*

*Table S3. Clinical studies for intramyocardial injections.*

*Table S4. Demographics of 5 human participants.*

### Materials and Methods

Study design

The primary objective of this study was to develop a multi-needle epicardial injector that can rapidly deliver therapeutics consistently into the entire left ventricular myocardium of a beating heart during open heart surgery. We hypothesized that our device could demonstrate the limitations of current methods explored in the clinic to deliver macromolecules into the myocardium such as epicardial patches, single bolus, and random multiple intramyocardial injections. We characterized diffusive and convective mass transport within the myocardium with *ex vivo* porcine myocardium. The effects of injection parameters such as needle depth, injection volume, flowrate, and needle insertion angle on injectate distribution were tested using *ex vivo* porcine myocardium. De-identified, diseased *ex vivo* human myocardium tissue was used to investigate the effects of collagen buildup on fluid flow in the myocardium. *Ex vivo* porcine myocardium was used to compare volumes of distribution of phosphotungstic acid following administration of an epicardial patch, single bolus injection, and multiple injections in an array. For fenestrated needles, hole parameters were obtained from simulations in COMSOL. Then, they were tested with *ex vivo* porcine myocardium to demonstrate injections through all three holes. Mechanical characterization of RISE-LV was performed with *ex vivo* porcine hearts. Demonstration of macromolecular delivery to the entire left ventricular myocardium and comparing total flow dimensions from single and multiple injections was performed in vivo in swine. All swine were euthanized 1 hour following drug injections in accordance with the animal protocols.

All procedures involving de-identified human tissue were reviewed by Institutional Review Boards (IRBs) of both Georgia Institute of Technology and Emory University. The study was approved by the Emory University IRB (STUDY00008066) and deemed a non-human study from the Georgia Tech IRB (Protocol H24231). Written consent for the study was waived from all subjects due to the use of de-identified tissue that were derived from tissue removed during an orthotopic heart transplantation surgery, for reasons independent of this study.

All animal study procedures were reviewed and approved by Emory University's Institutional Animal Care and Use Committee (STUDY IPROTO202400000099).

Each figure legend includes sample sizes for each experimental group. Sample sizes were determined based on similar studies in the literature. No data or outliers were excluded from any of the analyses.

Macromolecular distribution in *ex vivo*, healthy porcine myocardium

We identified flowrates that resulted in minimal backflow across a range of viscosities. Aqueous polyethylene glycol (3350 g/mol; Fisher Scientific, MA) solutions with varying mass fractions (0-0.5, 0.1 increments) were prepared, and their viscosities were measured with a rheometer (MCR302; Anton Paar, Austria). 60 $\mu$L of each solution was injected into the mid-wall of *ex vivo*, healthy adult porcine myocardium (3-5" diameter aortas; Cyrgus Company, NE) with a 27G hypodermic needle at varying flowrates (60-1200 $\mu$L/min) using a syringe pump (OEM Fusion 4000X Syringe Pump; ChemyX, TX). The volume of injected fluid that leaked out was calculated by dividing the mass change before and after injections by the density of the solution (n = 3).

To characterize macromolecular distribution in *ex vivo,* healthy porcine myocardium, we used 40 kDa fluorescein isothiocyanate-dextran (FITC-dextran; Milipore Sigma, Germany) as a proxy molecule of a $VEGF_{165}$ dimer (38 kDa). To simulate penetration by diffusion into the myocardium, 60 $\mu$L of a 3 mg/mL aqueous FITC-dextran solution was placed within a 5 mm radius cylinder attached onto the left ventricular epicardium with Loctite superglue (Loctite, CT) (fig. S11; n = 5). The same solution was injected into the mid-wall of the left ventricular myocardium with a 27G hypodermic needle inserted perpendicular to the epicardium at a flowrate of 60 $\mu$L/min with a syringe pump (n = 5). Histology and fluorescent microscopy were used for visualization of the injectate distribution (fig. S12). To compare the penetration depths from diffusion and convection, a two-tailed Student's *t*-test was performed for statistical significance.

To verify our hypothesis that injected fluids flow along myocardial fibers, we compared the flow angle of samples where the needles were inserted into the mid-wall (5-10 mm depth; n = 33) or near the epicardium (3 mm depth; n = 7). All samples were obtained by injecting 120 $\mu$L of 3 mg/mL FITC-dextran at 60 $\mu$L/min with a 27G hypodermic needle, perpendicular to the epicardium. These angles were calculated by measuring the change in distance between the center of the injectate and the right tissue boundary with respect to fluid flow in histological sections (see fig. S13 for schematics). A two-tailed Student's *t*-test was performed for statistical significance.

We quantified the effects of injectate volume, flowrate, and needle angle on fluid length, average height, and average width. Only one parameter was changed from the baseline of comparison (60 $\mu$L volume, 60 $\mu$L/min flowrate, and a 27G needle inserted perpendicular to the epicardium) each time. The following parameters were tested: injectate volume (60, 90, 120, and 240 $\mu$L; n = 5 for each volume); flowrate (60, 90, 120 $\mu$L/min; n = 5 for each flowrate); needle angle (90°, 45° with respect to the epicardium; n = 5 for each angle). Histology and fluorescent microscopy were used for visualization of the injectate distribution. For injectate volume and flowrate, a one-way analysis of variance (ANOVA) followed by Tukey's test was performed. For needle angle and size, a two-tailed Student's t-test was performed.

<u>Macromolecular distribution in ex vivo, diseased human myocardium</u>

All procedures were reviewed by Institutional Review Boards (IRBs) of both Georgia Institute of Technology and Emory University. The study was approved by the Emory University IRB (STUDY00008066) and deemed a non-human study from the Georgia Tech IRB (Protocol H24231). Written consent for the study was waived from all subjects due to the use of de-identified tissue that were derived from tissue removed during an orthotopic heart transplantation surgery, for reasons independent of this study. No outliers were excluded from the data. De-identified, diseased left ventricular myocardium tissue from 5 patients (table S4) were injected with the same solution (3 mg/mL 40 kDa FITC-dextran, aqueous) and injection parameters (120 $\mu$L, 60 $\mu$L/min, 27 G hypodermic needle, mid-wall). Histology and fluorescent microscopy were used for visualization of the injectate distribution. Picrosirius red staining was performed to qualitatively compare collagen content within the myocardium between diseased human tissue samples and healthy porcine myocardium samples.

Injectable hydrogel delivery to *ex vivo,* healthy porcine myocardium

An injectable hydrogel developed by Cohen *et al. (27)* was prepared and mixed with phosphotungstic acid (12.5% w/w), a contrast agent to model macromolecular transport in the myocardium. To simulate an epicardial hydrogel, excess crosslinker (calcium gluconate) was added to 540 $\mu$L of the hydrogel formulation and attached onto the epicardium of an *ex vivo,* healthy porcine heart (n = 3). For the single injection group, 540 $\mu$L of the hydrogel formulation was injected with a syringe pump at 60 $\mu$L/min into the mid-wall of the left ventricular myocardium with a 27G hypodermic needle (n = 3). For the multiple injections group, nine 60 $\mu$L injections were simultaneously performed in a 3 x 3 array (2 cm apart horizontally, 0.5 cm apart vertically) into the mid-wall with a prototype of RISE-LV (n = 3). 24 hours after the administration of the hydrogel, a microcomputed tomography (micro-CT) scan of the heart was performed to obtain the distribution of phosphotungstic acid.

Micro-CT contrast agent delivery using fenestrated needles

To enable injections targeting multiple layers of the myocardium, we sought to incorporate additional holes in 27G hypodermic needles. To identify hole size and location that enables equal flowrates out of all holes, we developed a COMSOL Multiphysics 6.3 (Sweden) model to simulate fluid flow from a fenestrated needle into the myocardium. The laminar flow physics model was used for flow within the needle. The injection flowrate was used for the inlet velocity conditions. For flow within the myocardium, the Darcy's law physics model was chosen. Porosity of 0.4 and permeability of $10^{-15}$ and $10^{-17}$ $m^2$ in the x- and y-directions, respectively, were chosen such that the modeled flow matched *ex vivo* results. A time-dependent simulation was performed to visualize injectate distribution during the injections. The no-slip boundary condition was used at all solid-fluid interfaces. Stagnant flow conditions were used for initial velocities, and an intra-tissue pressure of 2 atm was used.

Fenestrated needles with an additional 3 mm diameter circular hole placed 4 mm above the original bevel tip were fabricated with an OPTEC WS-Flex USP system (mark speed 50 mm/s, jump speed 100 mm/s, laser power 100%, burst time 1000 $\mu$s, laser on delay 100 $\mu$s, laser off delay 50 $\mu$s). With our syringe pump, we injected 240 $\mu$L of a 3 mg/mL phosphotungstic acid aqueous solution into the myocardium at 120 $\mu$L/min to account for the greater number of holes. We used micro-CT to visualize the distribution of phosphotungstic acid following the injection.

Design, fabrication, and mechanical characterization of RISE-LV

SolidWorks 2024 (Dassault Systèmes, France) was used to design components and molds for RISE-LV. The 5 mm thick backing contains 26-40 holes in an array, depending on the size of the heart; holes are placed 2 cm apart horizontally and 0.5 cm apart vertically. The backing was 3D-printed with a Form 3B (Formlabs, MA) resin printer with a biocompatible resin (BioMed Clear, Formlabs), washed in 99 % isopropyl alcohol (Florida Laboratories, FL) for 20 minutes, and cured with a Form Cure (Formlabs) at 60 ºC for 60 minutes. Elastomer molds were 3D-printed with a Bambu Lab X1C Printer (Bambu Lab USA, TX) with polylactic acid (PLA) filament (Bambu Lab USA). Uncured, equal masses of Ecoflex Part A and B were poured into molds to fabricate different elastomer layers and straps of RISE-LV. Additional uncured Ecoflex mixtures were poured onto the backing to embed the needles in Ecoflex, and to attach the needles onto the backing. Unless specified otherwise, Ecoflex was cured at room temperate for 5 hours. Then, a 0.5 cm-thick Ecoflex layer was attached under the backing with a small amount of uncured Ecoflex. For the elastomeric straps, Ecoflex was molded into two straps (2 cm width, 0.5 cm thick, 8 cm long; 2 cm width, 0.5 cm height, 3 cm long), onto which velcro was attached with a small amount of uncured Ecoflex. These straps were attached to the backing with partially cured Ecoflex (see fig. S14 for schematics).

For mechanical characterization of RISE-LV, pull tests were performed, wherein a mechanical test stand (Mark-10 ESM 750; Mark-10, NY) was used to pull on different components of RISE-LV at 1 cm/s unless specified otherwise. This speed translates to the approximate rate at which the heart contracts and relaxes at 60 beats per minute under anesthesia. The frictional force required to displace a single needle from the elastomer layers was measured by pulling on individual needles with the Mark-10 (n = 5). The force required to push all needles attached to RISE-LV into the myocardium was measured by pushing our device onto the heart with the Mark-10 (n = 3). To calculate the maximum pressure applied by the strap of RISE-LV onto the heart at various strain values, the strap was stretched from $\varepsilon = 0$ to $\varepsilon = 1$ for 20 cycles with the Mark-10 (n = 3). To calculate the force to displace RISE-LV from the heart, a modified prototype was fabricated with a customized backing with a handle for the force gauge grips. After strapping the modified prototype onto the heart such that the strain of the strap was 0.5, the backing was lifted with the Mark-10 (n = 3).

All *ex vivo* experiments with Ecoflex were repeated with Silopren™ LSR 2003 (LSR 2003; Momentive Performance Materials Inc., NY) with backings and molds 3D printed with High Temp V2 resin (Formlabs). Uncured, equal masses of LSR 2003 Part A and B were mixed and cured at 200 ºC for 10 minutes in an oven. Components 3D printed with High Temp V2 resin were washed in 99 % isopropyl alcohol for 5 minutes and cured with a Form Cure at 80 ºC for 120 minutes. For the strap, two straps were strapped around the heart such that the strain of each strap was 0.05.

Design and fabrication of a 20 syringe-rack for simultaneous injections

Solidworks 2024 was used to design a custom syringe rack capable of holding 20 1 mL syringes in a 10 x 2 array. Racks were 3D-printed with a Bambu Lab X1C 3D Printer with PLA filament. To simultaneously push all plungers, a 3/8 inch-thick aluminum block (McMaster-Carr, GA) was cut to match the profile of the syringe rack using a waterjet (OMAX Maxiem 1515; OMAX, WA). Our syringe pump was used to dispense water simultaneously from all 20 syringes with the 3D-printed rack and aluminum block. The volume of water dispensed from each syringe (n = 3) was calculated by dividing the mass differences of all centrifuge tubes before and after dispensing by the density of water and was compared to the programmed dispensed volume of the pump (120 μL).

mRNA-encapsulating lipid nanoparticle formulation

tdTomato mRNA was prepared as described previously (*36, 37*). The sequence of tdTomato was obtained as a gBlock from Integrated DNA Technologies (IA, USA). Using a three-molar excess of insert, the gBlock was cloned into a PCR amplified pMA7 vector. The resulting transcripts were purified using a 0.8% agarose gel and verified by Sanger sequencing. In vitro transcription was carried out overnight at 37°C using HiScribe T7 (New England BioLabs, MA) according to the manufacturer's instructions. The template was then removed by DNase I treatment. To finalize the mRNA preparation, a Cap1 structure was added following denaturation at 65°C for 10 minutes, and the transcripts were enzymatically polyadenylated. The purified RNA products were analyzed by gel electrophoresis to confirm purity.

Lipid nanoparticles carrying tdTomato mRNA were formulated using the NanoAssemblr Ignite (Precision NanoSystems, Canada). SM-102 was purchased from Organix Inc (MA), and cholesterol, PEG lipids and helper lipids were purchased from Avanti Research (AL). Lipids used were SM-102, 1,2-distearoyl-sn-glycero-3-phosphocholine (DSPC), cholesterol, and DMG-PEG (2K), at moderna lipid and RNA ratios (fig. S15A). Aqueous (25mM acetate) and organic (100% ethanol) phases were mixed at a flowrate ratio of 3:1 and total flowrate of 12 mL/min. LNPs were flash-frozen with 10% sucrose and stored in -80°C.

Nanoparticles were dialyzed using 100 kDa centrifugal filters into 10 mM Tris, then sterile purified through 0.22 μm filter. The hydrodynamic diameter of the LNPs was measured using dynamic light scattering (DynaPro Plate Reader III, Wyatt) (fig. S15, B and C), and the encapsulation efficiency was measured using RiboGreen assay (Invitrogen). The final concentration of lipid nanoparticles was 127 μg/ml.

Macromolecular distribution with RISE-LV *in vivo* in a swine model

All procedures were reviewed and approved by Emory University's Institutional Animal Care and Use Committee (STUDY IPROTO202400000099). All *in vivo* studies were performed at Emory University on female Yorkshire swine weighing approximately 30-40 kg (n = 3; Oak Hill Genetics, IL). From the weight of the swine, a smaller heart was expected; as a result, a smaller prototype with 26-27 needles was used. In addition, to accommodate for increased backflow due to the heart beating and hydration state in the porcine model, we arranged needles 1.5 cm apart horizontally rather than 2 cm apart.

We anesthetized the swine with intramuscular injections of TKX (telazol 4.4 mg/kg, ketamine 2.2 mg/kg, and xylazine 2.2 mg/kg), and the swine were intubated and maintained on 1-3% isoflurane in oxygen. A sternotomy was performed to access the left ventricle.

To compare the volume of distribution of single injections and multiple injections of the same total volume, we injected 40 kDa FITC-dextran (3 mg/mL) into the left ventricular free wall with a smaller prototype of RISE-LV with 5 needles (fig. S9). A small volume of tissue marking dye (Cancer Diagnostics, Inc., NC) was mixed to identify the injection sites for histological analysis. For single injections, 480 μL of the mixture was injected at 60 μL/min. For multiple injections, 4 needles in a 2 x 2 array (1.5 cm apart horizontally and 0.5 cm apart vertically) were used to inject 120 μL each at 60 μL/min. To demonstrate that we could deliver macromolecules to the entire left ventricular myocardium, a prototype of RISE-LV with 26-27 needles (9 x 4 array; 1.5 cm apart horizontally and 0.5 cm apart vertically) was used to deliver 120 μL of phosphotungstic acid from each needle at a flowrate of 60 μL/min. To demonstrate the delivery of functional mRNA-encapsulating lipid nanoparticles, we injected 480 μL of tdTomato mRNA-encapsulating lipid nanoparticles at 60 μL/min with the aforementioned smaller prototype. 480 μL was chosen to maximize the amount of mRNA delivered. tdTomato expression was confirmed with immunohistochemistry staining. No significant signs of trauma were observed during the procedure. All induced irregular heart rhythms were successfully controlled.

Micro-CT was performed to obtain the distribution profile of phosphotungstic acid in the left ventricular free wall portion. Distribution of FITC-dextran was obtained by histology and fluorescent microscopy. Hematoxylin & Eosin (H&E) staining was performed in needle insertion sites and sites where no needle was inserted into to qualitatively assess tissue damage.

<u>Analysis of the distribution of FITC-dextran through histology and fluorescent microscopy</u>

*Ex vivo* tissue samples were analyzed through cryosectioning. 24 hours after injections, tissue samples were embedded in optimal cutting temperature compound (OCT Compound; Sakura Finetek, CA), frozen, and sectioned with a cryostat (10-20 μm thickness) every 1 mm. *In vivo* tissue samples were fixed in 10% neutral buffered formalin (VWR), dehydrated, embedded in paraffin, and sectioned with a microtome (4-6 μm thickness) every 2 mm.

Fluorescent microscopy was performed (BioTek Cytation 7; Agilent Technologies, CA; GFP filter, brightness 5, exposure 32 ms, and gain 24) to visualize the distribution of FITC-dextran in each section. Individual tissue images were thresholded based on brightness with ImageJ to separate FITC-dextran from surrounding tissue. Custom python code was used to calculate the average height and width of injectate from histological sections. Length was calculated from the distance between tissue sections in which FITC-dextran was found.

In the analysis of *in vivo* experiments, the total flow dimensions of the 120 x 4 and RISE-LV groups represent the sum of lengths and average widths of all the injections in arrays.

<u>tdTomato detection through immunohistochemistry staining</u>

Goat Anti-tdTomato monoclonal antibodies were purchased from OriGene Technologies, Inc. (AB8181-200). Rabbit anti-goat secondary antibodies were purchased from Thermo Fisher (Catalog number 31402). A 3,3'-diaminobenzidine horseradish peroxidase substrate kit from Vector Laboratories, Inc. (CA; catalog number SK-4100) was used. Prior to performing the immunohistochemistry stains, tissue samples were fixed in 10% neutral buffered formalin, dehydrated, embedded in paraffin, and sectioned with a microtome (4-6 $\mu$m thickness) every 2 mm. Standard protocols from manufacturers were used for the immunohistochemistry stains. Stained sections were imaged with brightfield microscopy using a slide scanner (BioTek Cytation 7).

Analysis of the distribution of phosphotungstic acid through micro-CT

*Ex vivo* tissue samples were scanned with an Inveon CT scanner (Siemens, Germany; 70 kV, 500 mA). *In vivo* tissue samples were scanned with a VivaCT 40 scanner (Scanco Medical, Switzerland; 70 kV, 500 mA). 3D reconstruction was performed with Horos software (Horos Project, MD) and volume of distribution calculations were performed with Dragonfly 3D World (Comet Technologies, Canada). For the *in vivo* studies, lengths and average widths of individual injectates were obtained by analyzing projections onto the axial plane with python code. Average height of individual injectates were obtained by randomly sampling 5 injectates and analyzing images with python code.

Statistical analysis

All statistical analyses were conducting using GraphPad Prism 10. All data was assumed to follow normal distributions. Comparison of two groups was performed by an unpaired two-tailed Student's *t* tests. Comparison of three or more groups was performed by a one-way analysis of variance followed by Tukey's multiple comparison tests. Statistical significance was defined as $P < 0.05$ for all tests. Each figure legend contains sample sizes for each experimental group.

Supplementary Figures

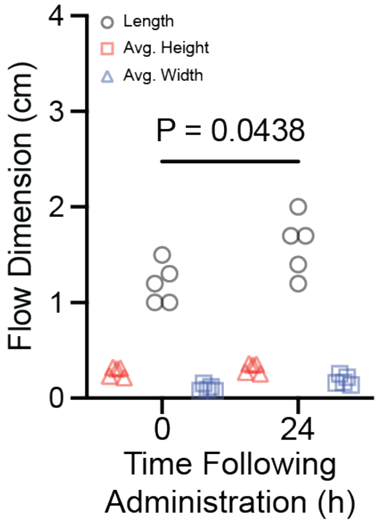


**Fig. S1. Injectate distribution after 0 and 24 hours.** Data from *ex vivo* healthy porcine left ventricular myocardium tissue samples; n = 5 tissue samples. Statistics: Two-tailed t test.

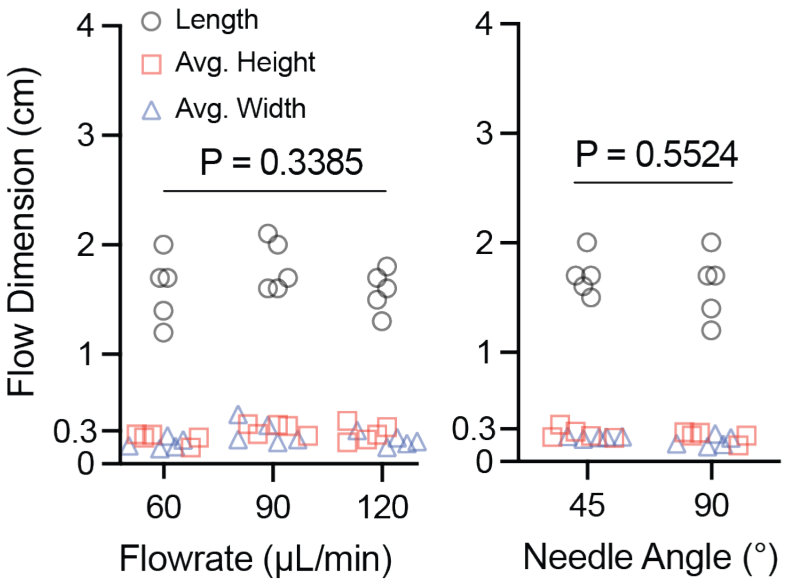


**Fig. S2. Effects of injection flowrate and needle angle on myocardial injectate distribution.** Data from *ex vivo* healthy porcine left ventricular myocardium tissue samples; n = 5 injections into different tissue samples for each group. Statistics: One way ANOVA followed by Tukey's multiple test for flowrate. Two-tailed t test for needle angle.

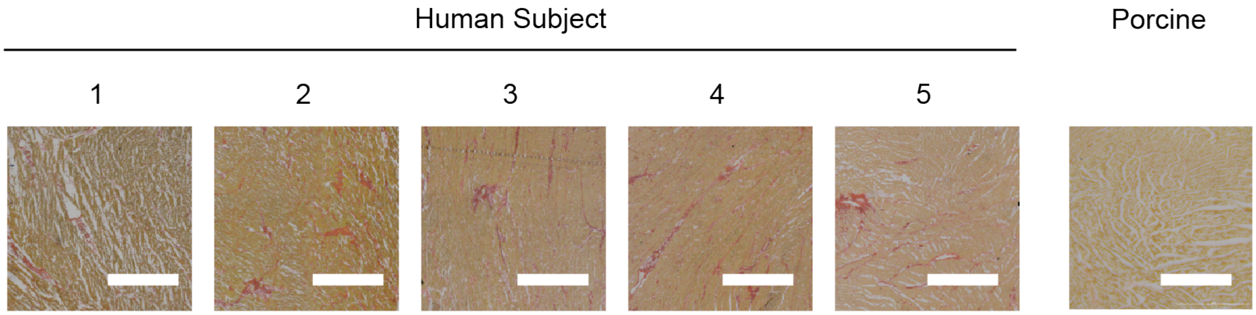


**Fig. S3. Histological stains of human and porcine myocardium samples.** Images of Picrosirius red-stained sections of the left ventricular myocardium. Muscle fibers and the cytoplasm is stained yellow while collagen is stained red. More collagen is seen in the diseased human tissue than the healthy swine tissue. Scale bar = 1 mm.

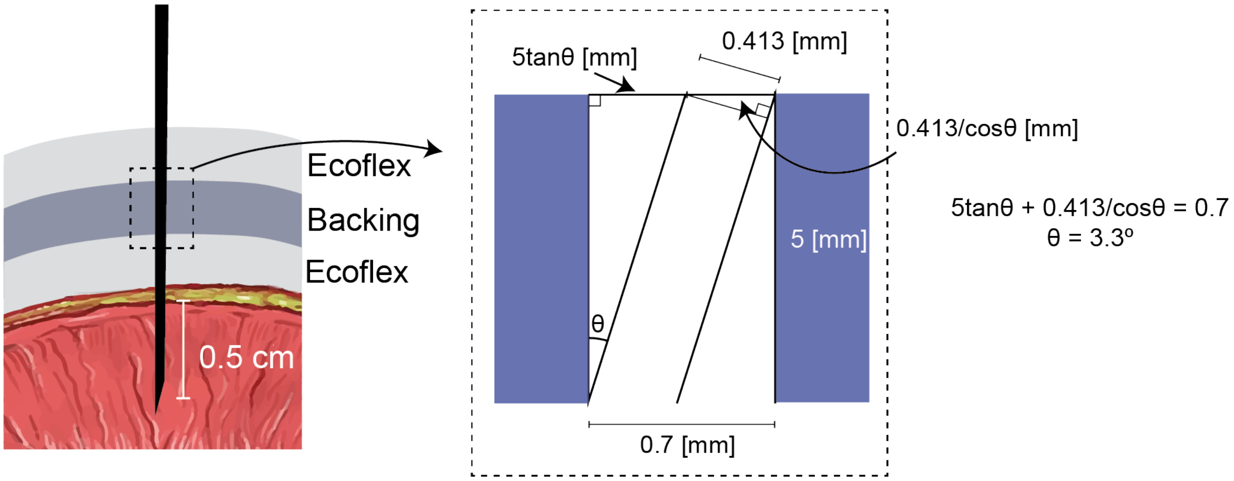


**Fig. S4. Design of the backing limiting vertical tilt angle deviation.** Illustration of the cross section of a hole in the backing and the calculated maximum vertical tilt angle, θ.

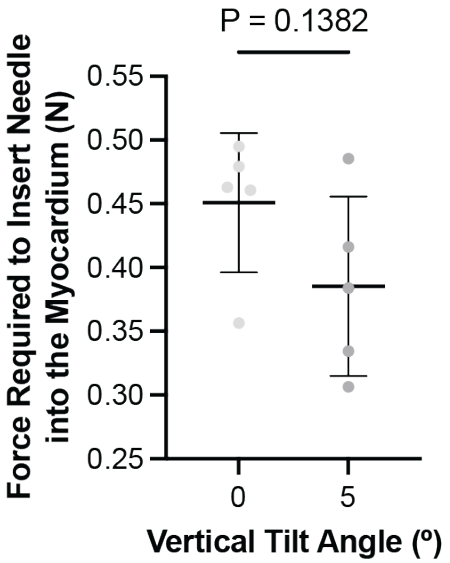


**Fig. S5. Single needle insertion force at 0º and 5º.** 0º represents minimum vertical tilt angle and 5º represents maximum vertical tilt angle. n = 5 needles in different swine myocardium tissue samples. Statistics: Two-tailed t test. Mean ± S.D.

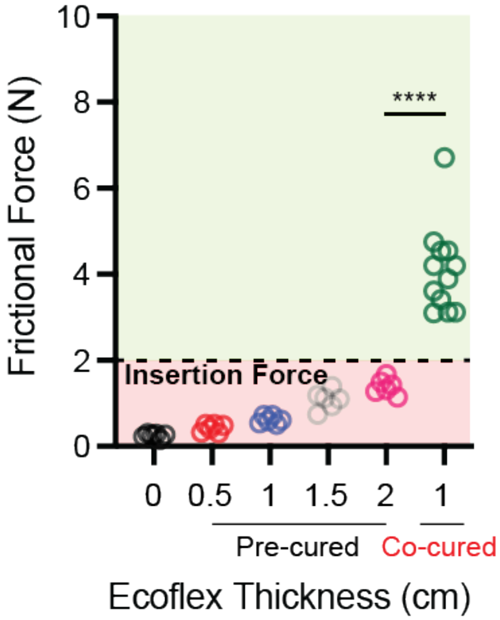


**Fig. S6. Single needle frictional force.** Frictional force provided by different thicknesses and curing methods of Ecoflex. $n = 5$-12 needles in different elastomer samples. Statistics: one way ANOVA followed by Tukey's multiple comparison test. ****$P < 0.0001$.

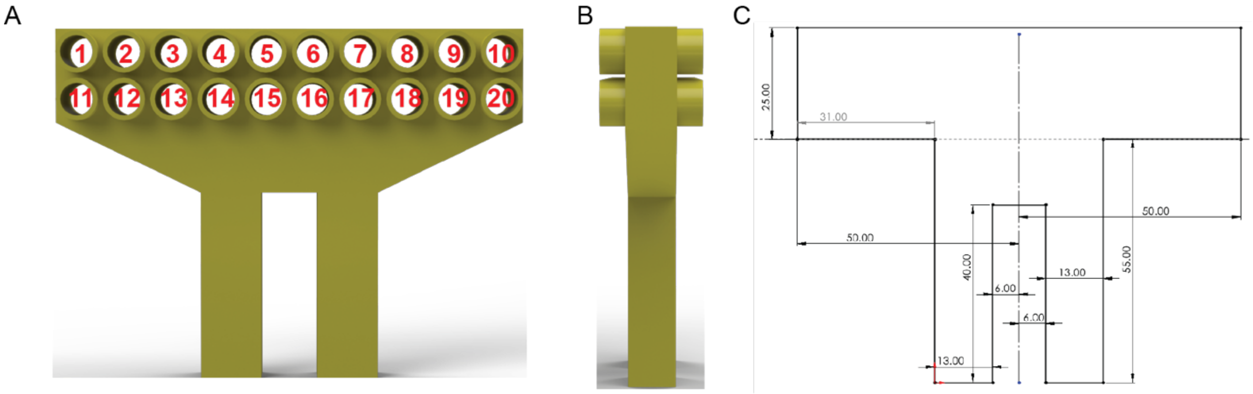


**Fig. S7. Syringe Rack Design & Characterization. (A**) Front view of a computer aided design of the syringe rack. Each position in the syringe rack is denoted by a number. (**B**) Side view of the syringe rack. (**C**) Dimensions of the aluminum block used to push all syringes simultaneously.

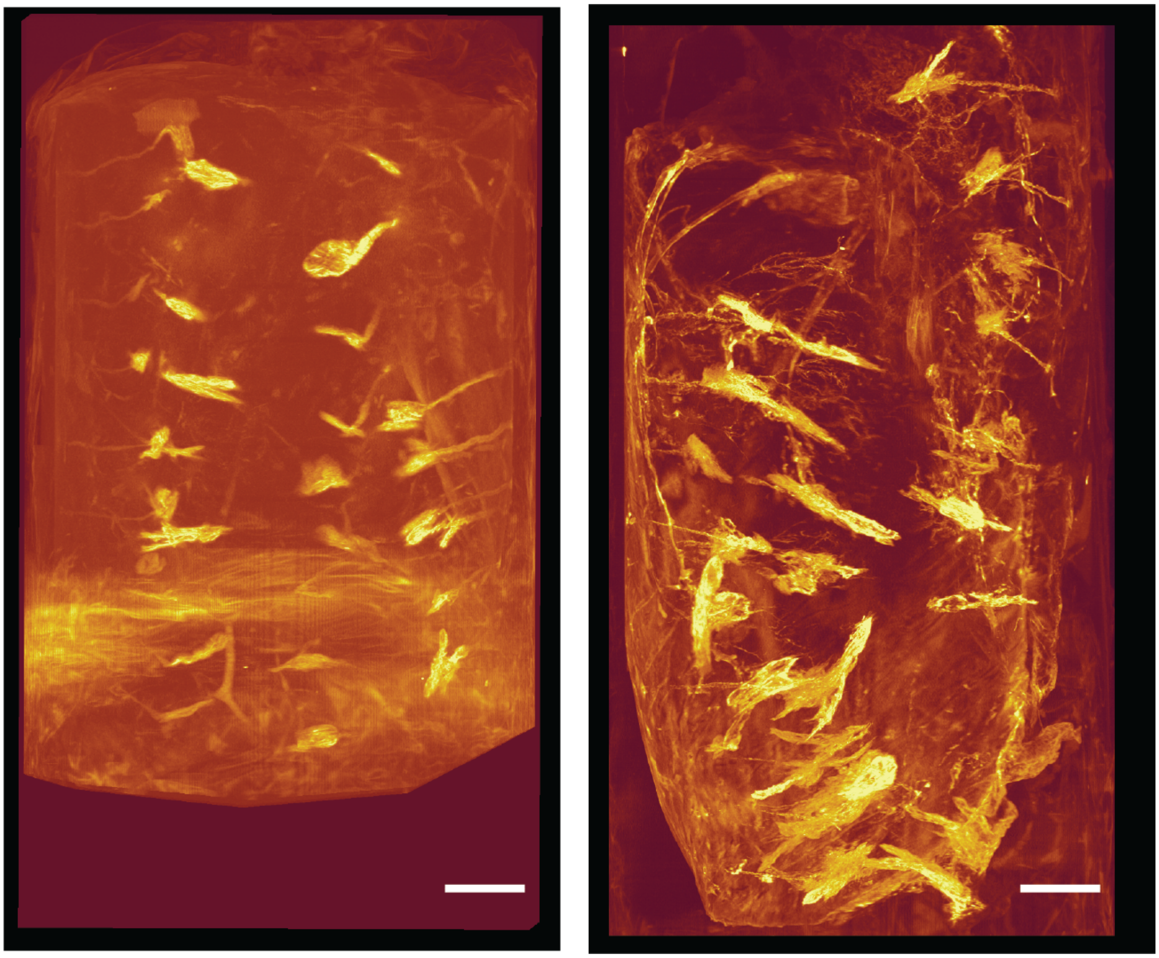

**Fig. S8. Micro-CT scans of the left ventricular myocardium of swine dosed *in vivo* with RISE-LV.** Bright yellow indicates phosphotungstic acid. Left: 2$^{nd}$ animal, right: 3$^{rd}$ animal. First animal micro-CT scan is found in Fig. 4. Scale bar = 1 cm.

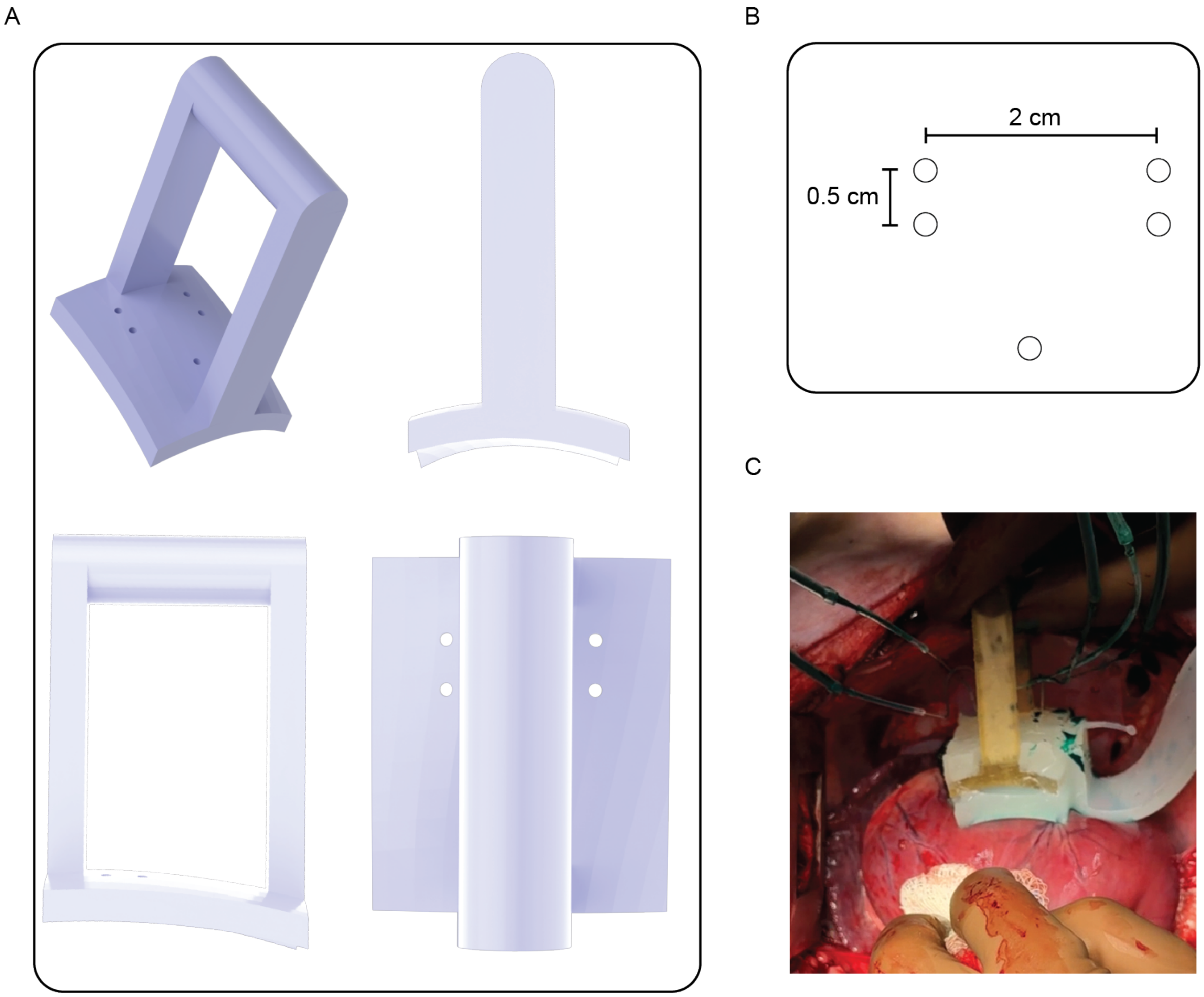


**Fig. S9. Four + one injection array RISE-LV used *in vivo* in some studies.** The four injection array allows for a small subsection of the myocardium to be injected with a solution. It also allows a comparison to a single injection of a 480 $\mu$L volume commonly used in clinical trials. (**A**) Clockwise from top left: isometric, front, top, and side view of the 3D computer aided design. (**B**) Location of holes in the smaller prototype. (**C**) Picture of a prototype being used to inject FITC-dextran and tissue marking dye *in vivo*.

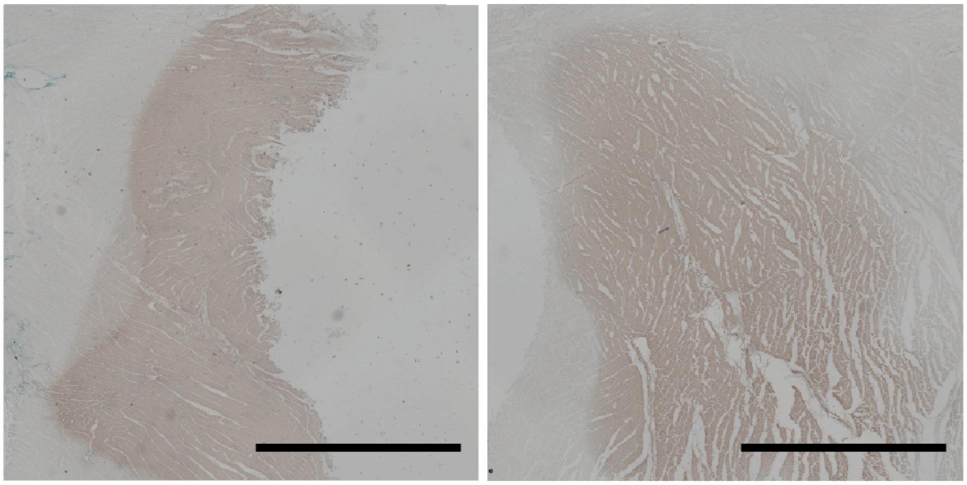

**Fig. S10. Representative Immunohistochemistry stains against tdTomato.** Brown indicates tdTomato. Left: 2nd animal, right: 3rd animal. Stain from the first animal is found in Fig. 4. Scale bar = 1 mm.

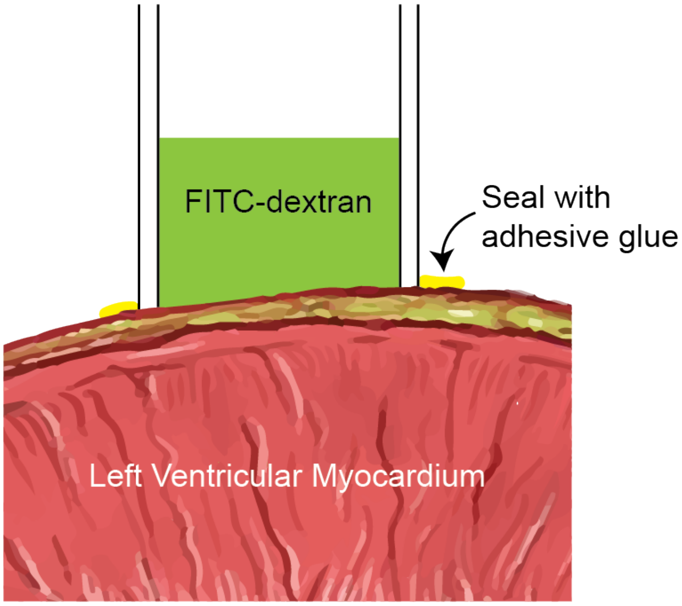


**Fig. S11. Setup of diffusion studies.** A 5 mm radius cylinder was attached to the left ventricular epicardium of *ex vivo* healthy porcine tissue using adhesive. 60 $\mu$L of FITC-dextran was added to the cylinder and allowed to diffuse into the tissue.

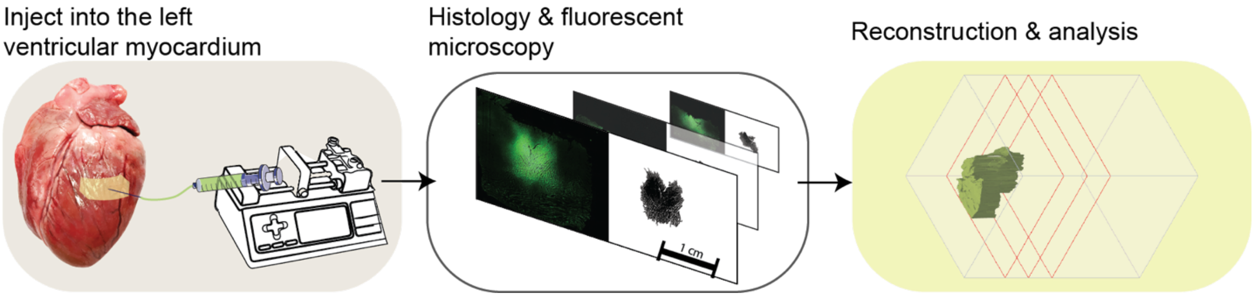


**Fig. S12. Injectate visualization by histology and microscopy.** Following injection of FITC-dextran, the myocardium is sectioned with a cryostat or microtome, and sections are imaged with a slide scanner. Images are thresholded based on brightness to determine the distribution of FITC-dextran, and reconstruction is performed with python code.

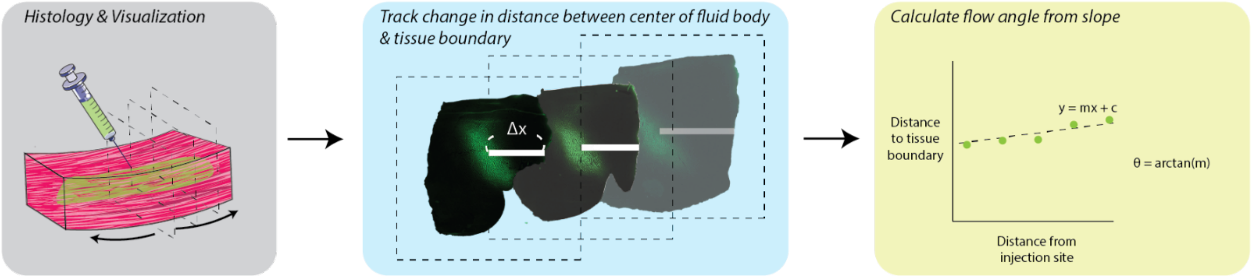


**Fig. S13. Flow angle calculation by histology and microscopy.** Following injection of FITC-dextran, the myocardium is sectioned with a cryostat or microtome and sections are imaged with a slide scanner. In each section, the distance between the center of the injectate and the right tissue boundary (Δx) is measured. The change in this value is used to calculate the flow angle.

1. Fabricate top & bottom Ecoflex layers by curing in a mold (60ºC, 2 hours).

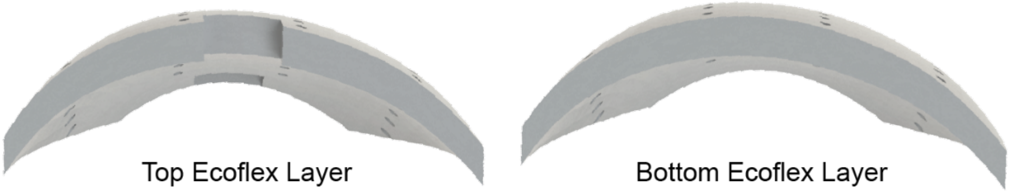


2. Attach top & bottom Ecoflex layers to the backing by applying a small amount of uncured Ecoflex onto the top & bottom surfaces of the backing. Cure at 60ºC for 1 hour.

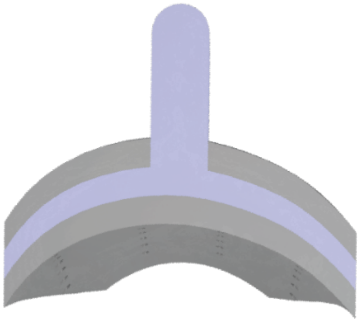

3. Fabricate elastomeric straps by curing in a mold (60ºC, 2 hours). The fabric texture represents Velcro. Velcro is placed on top of the elastomeric straps during the curing process.

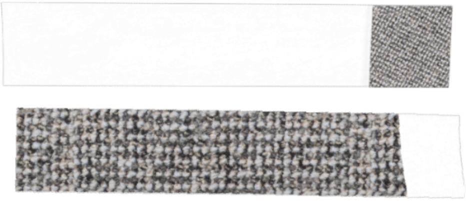

4. Attach straps to backing by applying a small amount of uncured Ecoflex onto the left and right surfaces of the backing. Cure at 60ºC for 1 hour.

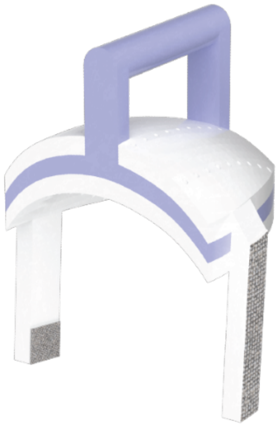

5. Insert needles into holes in the backing after placing the device onto the 2-part needle aligner (light green, light blue). The light green component contains 5 mm tall needle holes, and the light blue component is solid. This ensures that needles protrude no more or no less than 5 mm from the bottom Ecoflex layer.

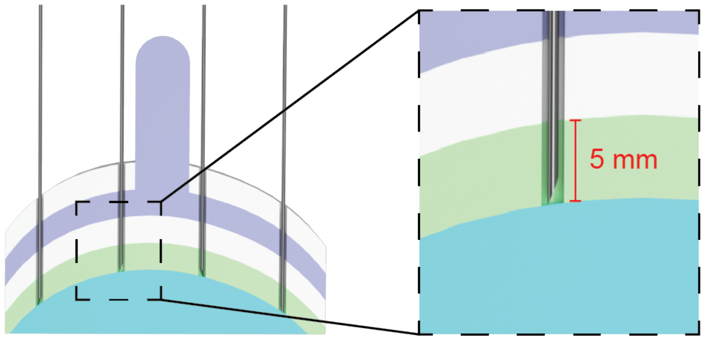


6. Attach needles onto backing and Ecoflex layers by pouring a small amount of uncured Ecoflex into needle holes.

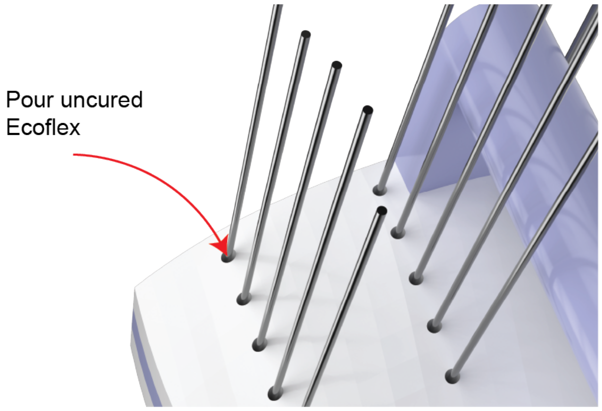


**Fig. S14. Procedure used to assemble RISE-LV prototypes.** Ecoflex layers (light grey) are fabricated by curing in a mold. These layers are attached to a backing (purple) with a small amount of uncured Ecoflex. Needle protrusion depths are controlled with hole depths in a 2-part aligner (light green and light blue). Needles are attached to the backing with a small amount of uncured Ecoflex.

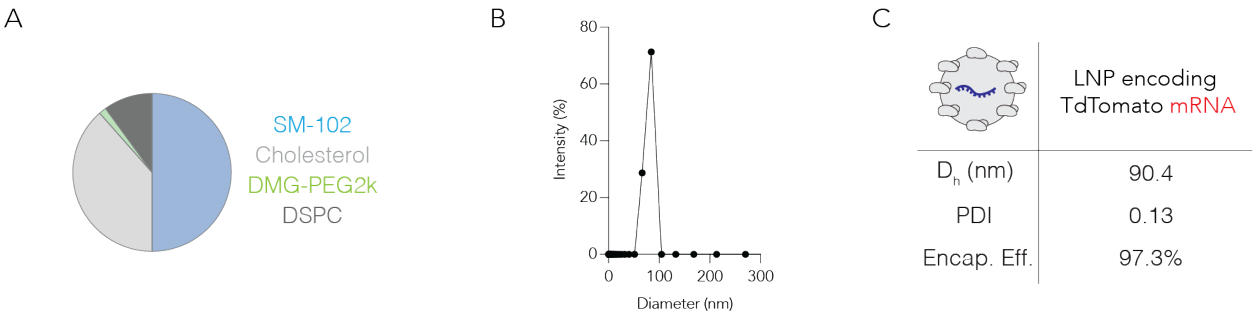


**Fig. S15. Formulation components and quality control of formulated tdTomato mRNA-encapsulating lipid nanoparticles.** (**A**) Lipid components of lipid nanoparticles. SM-102, cholesterol, DMG-PEG2000, and DSPC were combined at a molar ratio of 50:38.5:1.5:10 for formulation. (**B**) Sample dynamic light scattering readout with diameter and intensity. (**C**) Diameter ($D_h$; nm), polydispersity index (PDI), and encapsulation efficiency (%) of formulated and frozen tdTomato mRNA LNP following freeze-thaw.

<u>Supplementary Tables</u>

| *Swine Heart Delivery* | *Mass Transport Mechanism* | *Transported Species* | *Length Scale* | *Spacing* | *Administration Procedure* |
|---|---|---|---|---|---|
| *ex vivo* | Diffusion (24 h) | 40 kDa FITC-dextran | 0.2 cm (*h*) | N/A | Suture to epicardium |
| *ex vivo* | Convection (60 $\mu$L/min) | 40 kDa FITC-dextran | 1.2 cm (*l*) x 0.3 cm (*h*) x 0.1 cm (*w*) | N/A | Transepicardial Injection |
| *ex vivo* | Convection (60 $\mu$L/min) + Diffusion (24 h) | 40 kDa FITC-dextran | 1.6 cm (*l*) x 0.3 cm (*h*) x 0.3 cm (*w*) | N/A | Transepicardial Injection |
| *ex vivo* | Multi-Hole Convection (60 $\mu$L/min) + Diffusion (24 h) | Phosphotungstic acid | 1.6 cm (*l*) x 0.9 cm (*h*) x 0.3 cm (*w*) | 0.3 cm (*h*) | Transepicardial Injection |
| *in vivo* | **RISE-LV (60 $\mu$L/min)** | **40 kDa FITC-dextran** | **3.5 cm (*l*) x 0.2 cm (*h*) x 2.2 cm (*w*)** | **1.5 cm (*l*), 0.5 cm (*w*)** | **Transepicardial Multi-Injection** |

**Table S1. Summary of mass transport dimensions in porcine hearts.** RISE-LV optimizes total tissue coverage in the heart.

| *Device* | *Mass Transport Mechanism* | *Transported Species* | *Length Scale* | *Application Time* | *Transport Time* | *Spacing* | *Administration Procedure* | *Removal Procedure* | *Reference* |
|---|---|---|---|---|---|---|---|---|---|
| **RISE-LV (60 μL/min)** | **Convection** | **40 kDa FITC-dextran** | **3.5 cm (*l*) x 0.2 cm (*h*) x 2.2 cm (*w*)** | **mins** | **0.1 h** | **1.5 cm (*l*), 0.5 cm (*w*)** | **Transepicardial Multi-Injection** | **N/A** | |
| AAV-loaded Microneedle Array | Diffusion | Adenovirus | 0.1 cm (*h*) x 0.05 cm (*w*) | mins | Not reported | N/A | Microneedle Applicator | N/A | 2 |
| Cardiac Cell-loaded Microneedle Patch | Diffusion | Cardiac Stromal Cells | 0.03 cm (*h*) | wks | 72 h | N/A | Microneedle Patch + Fibrin Glue | Open Heart Surgery | 3 |
| Refillable Polymer Reservoir | Diffusion | 40 kDa FITC-dextran | 0.2 cm (*h*) | wks | 24 h | N/A | Suture to epicardium | Open Heart Surgery | 4 |

**Table S2. Preclinical device comparisons for cardiac drug delivery.** Many preclinical cardiac drug delivery devices are not optimized to maximize tissue coverage in large animal models.

| Study Name | Mass Transport Mechanism | Injection Site Determination | Volume /Injection ($\mu L$) | Number of Injections | Injection Flowrate | Spacing | Administration Procedure | Removal Procedure | Reference |
|---|---|---|---|---|---|---|---|---|---|
| NCT03370887 | Convection | [$^{15}$O]-water positron emission tomography | 200 | 30 | Manual (> 240 $\mu$L/min) | ~1 cm ($l$, $w$) | Transepicardial Injection | N/A | 20 |
| NCT01174095 | Convection | 99mTc-sestamibi myocardial perfusion scintigraphy | 100 | 10 | Manual (> 240 $\mu$L/min) | ~1-1.5 cm ($l$, $w$) | Transepicardial Injection | N/A | 11 |
| EudraCT 2005-003629-19 | Convection | Electromechanical mapping | 100 | 20 | Manual (> 240 $\mu$L/min) | N/A | Mapping Catheter - Transendocardial Injection | N/A | 21 |
| phVEGF2 Trial | Convection | Electromechanical mapping | 1000 | 6 | Manual (> 240 $\mu$L/min) | N/A | Mapping Catheter - Transendocardial Injection | N/A | 22 |
| Euroinject One phase II Trial | Convection | Electromechanical mapping | 300 | 10 | Manual (> 240 $\mu$L/min) | N/A | Mapping Catheter - Transendocardial Injection | N/A | 23 |
| KAT-301 | Convection | Electromechanical mapping | 200 | 10 | Manual (> 240 $\mu$L/min) | N/A | Mapping Catheter - Transendocardial Injection | N/A | 24 |

**Table S3. Clinical studies for intramyocardial injections.** Many clinical trials utilize different methods to determine injection sites and injection parameters different to the optimized parameters found in our study.

| Subject | Age | Sex | Race | Height (m) | Weight (kg) | BMI (kg/m$^2$) | Disease |
|---|---|---|---|---|---|---|---|
| 1 | 31 | M | B | 1.82 | 57 | 17.21 | Nonischemic cardiomyopathy |
| 2 | 57 | F | B | 1.49 | 65 | 29.28 | Nonischemic cardiomyopathy |
| 3 | 52 | F | B | 1.72 | 98 | 33.13 | Nonischemic cardiomyopathy |
| 4 | 40 | M | B | 1.78 | 88 | 27.77 | Nonischemic cardiomyopathy |
| 5 | 26 | F | B | 1.52 | 64 | 27.70 | Nonischemic cardiomyopathy |

**Table S4. Demographics of 5 human participants.** Height and weight of subjects measured prior to heart transplant surgeries. M: Male, F: Female, B: Black
